# Independent component analysis (ICA) applied to dynamic oxygen-enhanced MRI (OE-MRI) for robust functional lung imaging at 3 T

**DOI:** 10.1101/2023.07.05.547787

**Authors:** Sarah H. Needleman, Mina Kim, Jamie R. McClelland, Josephine H. Naish, Marta Tibiletti, James P. B. O’Connor, Geoff J. M. Parker

## Abstract

**Purpose:** Dynamic lung oxygen-enhanced MRI (OE-MRI) is challenging due to the presence of confounding signals and poor signal-to-noise ratio, particularly at 3 T. We have created a robust pipeline utilizing independent component analysis (ICA) to automatically extract the oxygen-induced signal change from confounding factors to improve the accuracy and sensitivity of lung OE-MRI.

**Methods:** Dynamic OE-MRI was performed on healthy participants using a dual-echo multi-slice spoiled gradient echo sequence at 3 T and cyclical gas delivery. ICA was applied to each echo within a thoracic mask. The ICA component relating to the oxygen-enhancement signal was automatically identified using correlation analysis. The oxygen-enhancement component was reconstructed, and the percentage signal enhancement (PSE) was calculated. The lung PSE of current smokers was compared with non-smokers; scan-rescan repeatability, ICA pipeline repeatability, and reproducibility between two vendors, were assessed.

**Results:** ICA successfully extracted a consistent oxygen-enhancement component for all participants. Lung tissue and oxygenated blood displayed opposite oxygen-induced signal enhancements. A significant difference in PSE was observed between the lungs of current smokers and non-smokers. The scan-rescan repeatability, and the ICA pipeline repeatability, were good.

**Conclusion:** The developed pipeline demonstrated sensitivity to the signal enhancements of the lung tissue and oxygenated blood at 3 T. The difference in lung PSE between current smokers and non-smokers indicates a likely sensitivity to lung function alterations that may be seen in mild pathology, supporting future use of our methods in patient studies.

## 1. Introduction

Dynamic oxygen-enhanced MRI (OE-MRI) is a functional imaging technique that utilizes inhaled oxygen as a contrast agent.^1, 2^ When applied to the lung, observation of the spatial distribution and temporal evolution of the OE-MRI signal change can provide information relating to the delivery and uptake of oxygen.^3^ However, analysis of dynamic lung OE-MRI is challenging due to the presence of confounds including: artefacts and proton density changes arising from cardiac and respiratory motion; artefacts due to blood flow; poor signal-to-noise ratio of lung tissue resulting from the extremely short parenchymal T * and its low proton density. The impact of these confounds masks the small amplitude oxygen-induced signal change, which can reduce the sensitivity and accuracy of lung OE-MRI.

Most lung OE-MRI studies have focused on measuring T_1_-related signal enhancements at 1.5 T. As 3 T scanners are routinely used in clinical settings, development of lung OE-MRI at this field strength is important to enable future clinical translation. However, as the MR field strength is increased, the T * dephasing increases^4^ and the longitudinal relaxivity of oxygen decreases.^5^ These can result in a greater masking of the underlying OE-MRI signal by confounding factors and can reduce the accuracy and T_1_ sensitivity at 3 T relative to 1.5 T. Due to the greater impact of the confounding factors on lung OE-MRI at 3 T, it is particularly attractive to develop approaches to separate the oxygen-induced signal from confounds at this field strength.

Independent component analysis (ICA) is a data-driven blind source separation technique for extracting different signal sources from measured data. The extracted signal sources, known as ICA components, linearly combine to form the measured data. The ICA components are assumed to be independent of each other.^6, 7^ Isolation of the oxygen- enhancement signal response using ICA was demonstrated by Moosvi et al. in pre-clinical tumors.^8^ In this paper, we employ ICA to extract the oxygen-enhancement signal from confounds to improve the sensitivity of OE-MRI to alterations in lung function. We demonstrate the method in an experiment comparing the oxygen-induced signal changes seen in healthy smokers and non-smokers. The scan-rescan repeatability, the ICA pipeline repeatability, and the multi-site multi-vendor reproducibility, of the technique are investigated.

## 2. Theory

Several different MR contrast mechanisms are induced in lung OE-MRI, which contribute to the measured oxygen-enhancement response. During the inhalation of pure oxygen, excess molecular oxygen dissolves in lung tissue water and oxygenated pulmonary capillaries and veins. Molecular oxygen is paramagnetic, hence an increased concentration of dissolved oxygen causes T_1_ shortening in lung tissue and oxygenated blood^1^ (illustrated by contrast mechanism A in Figure 1).

**Figure 1:**
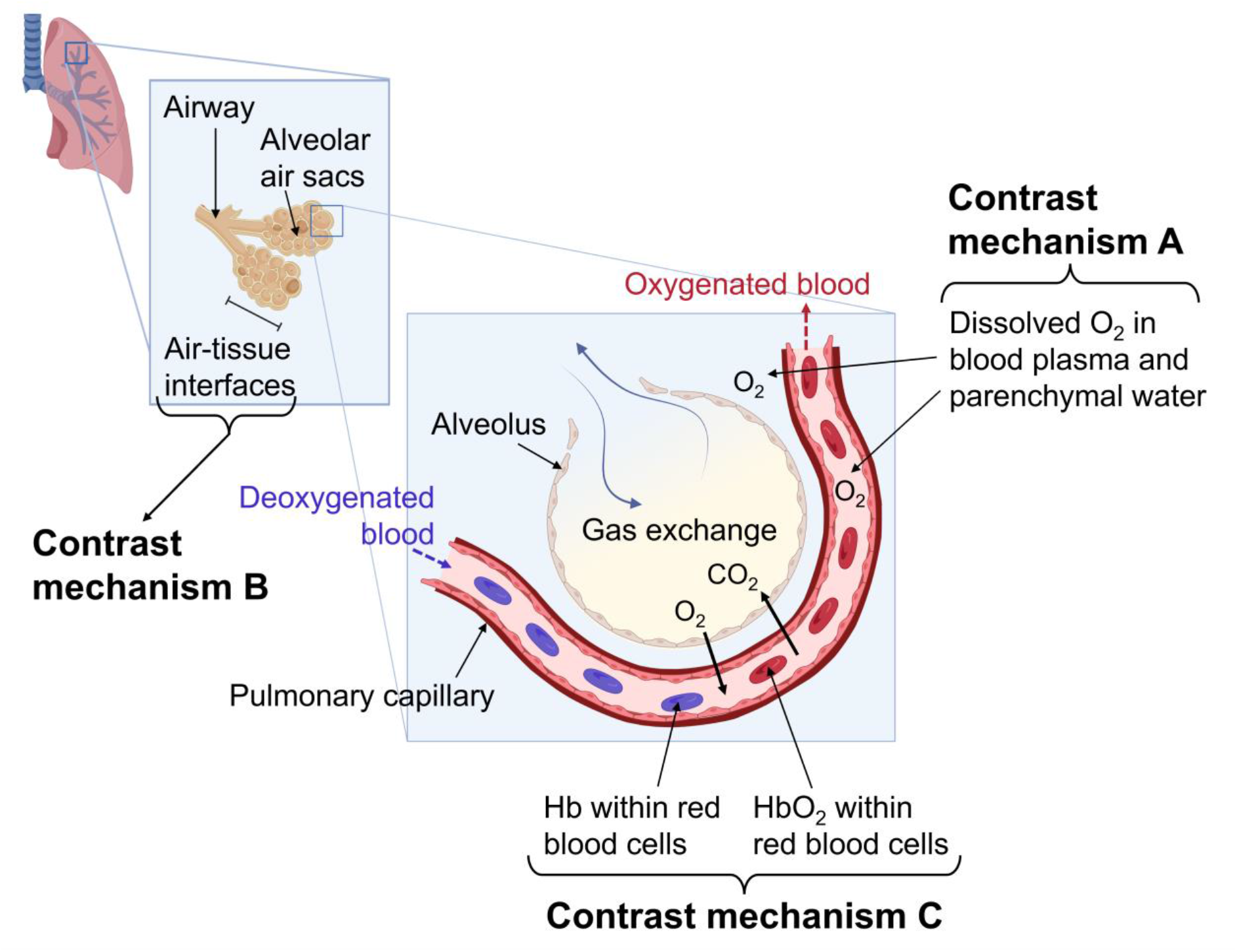
Diagram illustrating the main oxygen-induced contrast mechanisms of the lung. **Contrast mechanism A:** During air-inhalation, the blood is near full oxygen (O_2_) saturation in healthy participants. During 100% O_2_-inhalation, excess O_2_ dissolves in blood plasma and lung tissue water. Dissolved O_2_ is paramagnetic, which can shorten T_1_. Hence, the increased concentration of dissolved O_2_ during 100% O_2_-inhalation causes T_1_ shortening of lung tissue and oxygenated blood. **Contrast mechanism B:** Magnetic susceptibility gradients arise due to gas-tissue interfaces in the lung, which can shorten T *. Gaseous O has a greater magnetic susceptibility than air. As a consequence, 100% O_2_-inhalation increases the lung susceptibility gradients and causes T * shortening of lung tissue. **Contrast mechanism C:** Deoxygenated hemoglobin molecules (Hb) contained within red blood cells are paramagnetic. The paramagnetic Hb molecules generate magnetic susceptibility distortions which can shorten T *. In the blood, O_2_ binds to Hb molecules to form oxyhemoglobin (HbO_2_). However, Hb molecules in the blood are close to full O_2_ saturation during air-inhalation. 100% O_2_-inhalation therefore results in a minor increase in blood oxygen saturation. The Hb concentration decreases, causing the T * of oxygenated blood to increase (BOLD contrast). Created with BioRender.com.

Figure 1 also illustrates the two major T * contrast mechanisms in the lung. Represented by contrast mechanism B, T * can be affected by changes in magnetic susceptibility arising due to gas-tissue interfaces in the lung. As gaseous oxygen has a greater magnetic susceptibility than air, the inhalation of pure oxygen increases the susceptibility gradients within the lung and causes T * shortening.^9, 10^

Contrast mechanism C demonstrates the effect of deoxyhemoglobin on T *. Deoxyhemoglobin is paramagnetic and creates local magnetic field distortions that induce T * shortening (i.e. BOLD contrast).^11^ Oxygenated pulmonary blood is close to full saturation during air-inhalation for healthy participants.^12^ Consequently, the inhalation of pure oxygen results in a minimal change to the blood oxygen saturation, producing a small BOLD effect.^13^

## 3. Methods

### 3.1. Single Center OE-MRI acquisition

23 healthy participants were imaged on a 3 T Philips Ingenia MRI scanner after local institutional review board approval (18837/001, UCL Research Ethics Committee) and written informed consent. The recruited participants had no previous record of lung disease; participant information is summarized in Table 1(A). The dynamic OE-MRI protocol used a 2D coronal multi-slice dual-echo T1-fast field echo (RF-spoiled gradient echo) sequence to acquire four posterior slices during free-breathing.^14^ Echo times of TE_1_/TE_2_ = 0.72/1.2 ms were used; full sequence details are provided in Supporting Information Table S2(A).

**Table 1:**
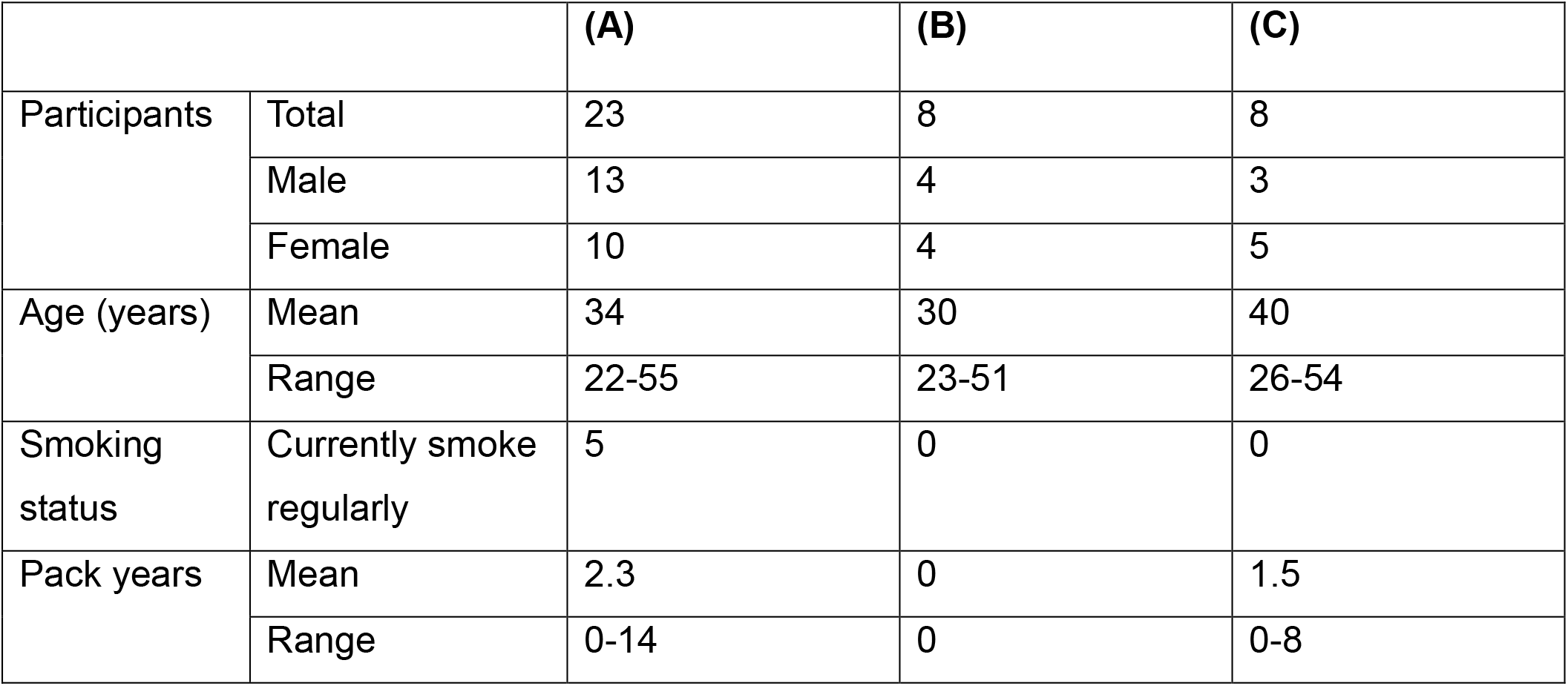
Summary of participant information for (A) the healthy participant study, (B) the repeatability study and (C) the reproducibility study. All recruited participants were healthy and had no previous record of lung disease. Some participants were common across groups. Further details of the non-smokers and current smokers involved in the healthy participant study (A) are provided in Supporting Information Table S1.

|  |  | (A) | (B) | (C) |
| --- | --- | --- | --- | --- |
| Participants | Total | 23 | 8 | 8 |
|  | Male | 13 | 4 | 3 |
|  | Female | 10 | 4 | 5 |
| Age (years) | Mean | 34 | 30 | 40 |
|  | Range | 22-55 | 23-51 | 26-54 |
| Smoking status | Currently smoke regularly | 5 | 0 | 0 |
| Pack years | Mean | 2.3 | 0 | 1.5 |
|  | Range | 0-14 | 0 | 0-8 |

Participants inhaled medical air (21% O_2_) and pure oxygen (100% O_2_) during the dynamic scan; gas was delivered via a non-rebreathing face mask (Intersurgical Ltd., Wokingham, UK) at a flow rate of 15 L min^-1^. The delivered gas was cycled between air and 100% O_2_ three times as depicted in Supporting Information Figure S1. The cyclic gas delivery imposed a temporal modulation on the oxygen-enhancement MR signal, designed to increase ICA sensitivity^8^ and aid identification of the ICA component relating to the oxygen- enhancement response. The delivered gas was manually switched between air and 100% O_2_ every 1.5 minutes using a gas blender (IHC Low Flow Blender, Inspiration Healthcare Ltd., Earl Shilton, UK). The 1.5-minute gas period was chosen to minimize the total dynamic scan duration, thereby maximizing participant scan tolerability in future patient studies.

### 3.2. Data post-processing

Motion correction of the dynamic images was performed using the deformable image registration software NiftyReg.^15^ The motion correction parameters implemented in NiftyReg are provided in Supporting Information Table S3. The 2D motion correction aligned all dynamic images with a reference image, processed separately for each slice. The reference image was chosen to represent the mean lung position and was identified as the dynamic image with the greatest correlation to the average of all the images in the dynamic series.

The reference images were manually segmented by intensity thresholding in ImageJ.^16^ Two anatomical masks were created (shown in Supporting Information Figure S2): a mask of lung tissue excluding major vasculature, and a mask of the full thoracic cavity including the heart and major vessels.

### 3.3. Application of ICA

Temporal ICA was applied to the thoracic masked dynamic OE-MRI time series using scikit-learn (version 1.2.2) FastICA,^17, 18^ separately for each echo. The number of ICA components separated during the analysis can affect the form of the resulting components. However, determination of the optimum number of components to separate is an unsolved problem and the optimum number of components may vary between subjects. Additionally, the ordering of the ICA components is arbitrary. Therefore, it was necessary to identify the ICA component relating to the oxygen-enhancement signal (referred to as the “OE ICA component”) and the number of ICA components to use for each dataset.

ICA was repeatedly run using an increasing number of components from 22 to 72 (51 separate instances) for each dataset. Initial experiments showed this range of component numbers enabled the OE ICA component to be reliably identified and use of fewer or more components did not improve the ability of ICA to extract the OE ICA component. The Spearman correlation coefficient of a sinusoidal approximation of the oxygen-induced signal (Supporting Information Figure S3) to every component from all runs of ICA was calculated using SciPy.^19^ The correlation values were compared across all ICA components and the single component with the greatest correlation value was identified as the optimal OE ICA component for the dataset under consideration (illustrated in Supporting Information Figure S3). Hence, the ICA pipeline enables the automatic identification of the optimal OE ICA component and overcomes the ambiguity of the number of ICA components to use.

If the magnitude of the correlation coefficient of the identified optimal OE ICA component was less than 0.4, the ICA application process was repeated for the dataset under consideration. A correlation value of less than 0.4 indicated the extraction of a poor OE ICA component, likely due to the ICA algorithm getting stuck in a bad local optimum. Known as algorithmic uncertainty, the convergence of ICA at a local optimum, rather than the global optimum, may occur due to the random initialization of ICA.^20, 21^ The algorithmic uncertainty of ICA within the pipeline was investigated and is described later.

The MR signal originating from the OE ICA component was reconstructed for comparison with the raw motion-corrected MRI data. The percentage signal enhancement (PSE) of the reconstructed OE ICA component data (PSE_ICA_) and the raw motion-corrected MRI data (PSE_MRI_) were calculated. The PSE describes the MR signal change that occurred upon the inhalation of pure oxygen and can be used to map the oxygen-induced signal response across the lung.

The PSE time series were calculated using Equation (1) where SI_air_ is the mean air- inhalation image, and SI(*t*) is the dynamic image at time *t*. SI_air_ was calculated by averaging the first 60 dynamic images. PSE maps were generated using the mean air-inhalation image, SI_air_, and the mean oxygen-inhalation image, SI_oxy_, in Equation (2). SI_oxy_ was calculated by averaging the final five dynamic images from each of the three oxygen- inhalation periods (a total of 15 images). Hence, for the cyclic gas delivery scheme with 1.5- minute gas periods, the PSE maps describe the signal enhancement that occurred during 1.5 minutes of oxygen inhalation.

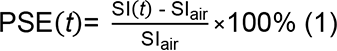

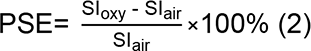

The median PSE_ICA_ map value within the lung mask was assessed for each subject. The median lung PSE_ICA_ values were compared between non-smoker and current smoker participants using an unpaired (independent samples) t-test; *p* < 0.05 were considered significant. An identical comparison was made for the median lung PSE_MRI_. Multiple regression was employed to adjust for the confounds of age and gender on the smoking status comparison. Separate models were created for each set of data (both echoes of PSE_ICA_ and PSE_MRI_) containing current smoking status, age, and gender, as variables. All statistical analyses were performed using IBM SPSS Version 28.0 (SPSS Statistics for Windows, IBM Corp.).

### 3.4. Repeatability and reproducibility

4-6 week (mean 5 weeks) repeat scans of 8 healthy participants were performed to assess the repeatability of the developed ICA OE-MRI analysis technique. The scan-rescan repeatability study used the 3 T Philips Ingenia scanner with protocol settings as described above. A reproducibility study was also carried out: 8 healthy participants were scanned on the same 3 T Philips Ingenia scanner and on a 3 T Siemens MAGNETOM Vida scanner located in a different institution using a multi-slice 2D double echo FLASH acquisition, within a 4-6-week interval (mean 5 weeks). Due to sequence implementation differences, echo times were increased to TE_1_/TE_2_ = 0.81/1.51 ms for the Siemens scan (further details of the Siemens implementation are provided in Supporting Information Table S2(B)). Participant information for the scan-rescan and reproducibility studies are summarized in Table 1 (B) and (C), respectively. One scan-rescan participant was excluded from the analysis due to the presence of substantial diaphragm ghosts in the second scan.

The repeatability and reproducibility scans were processed and analyzed using ICA as described above. For the scan-rescan repeatability study, the median lung PSE_ICA_ was compared between the repeat scans of each echo using Bland-Altman analysis including bias and limits of agreement (LoA), the repeatability coefficient (RC), and two-way single measure mixed-effects model intra-class correlation coefficient (ICC) with absolute agreement.^22, 23^ The ICC values were taken to indicate moderate repeatability if between 0.5- 0.75, good repeatability if between 0.75-0.9, and excellent repeatability if greater than 0.9.^24^ Variations in the measured PSE between the Siemens and Philips scans were expected due to the different echo times implemented (predicted by sequence-specific signal simulations^14, 25^ shown in Supporting Information Figure S4). As a result, we were unable to perform direct comparisons of PSE_ICA_ for the reproducibility study. Bland-Altman analysis was used to compare the echo time trend of the reproducibility study PSE_ICA_ to the signal simulations.

The repeatability of ICA within the pipeline was separately examined to assess algorithmic uncertainty by a repeat application of the pipeline to the scan-rescan participants. As for the scan-rescan study, Bland-Altman analysis was performed and the RC and ICC were calculated, to compare the median lung PSE_ICA_ between repeat pipeline applications.

## 4. Results

### 4.1. Extraction of the oxygen-enhancement signal response

The ICA analysis pipeline successfully extracted and identified the optimal OE ICA component from both echoes of every participant. The cyclic OE-MRI protocol was tolerated well by all participants and no adverse events were recorded. Across the participants, the optimal OE ICA component was found in runs of ICA in which different numbers of ICA components were used: range of 22-53 components (mean 30) for echo 1 and range of 22- 52 components (mean 30) for echo 2. The non-smoker PSE_ICA_ maps demonstrated consistent signal distributions within the lung (each subject presented in Figure 2(A)) with minor variations between maps due to the slice location, blood vessel content and presence of cardiac tissue. The PSE_ICA_ time series of all non-smokers displayed a cyclical signal enhancement in response to the air-oxygen gas switching (each subject presented in Supporting Information Figure S5(A)).

**Figure 2:**
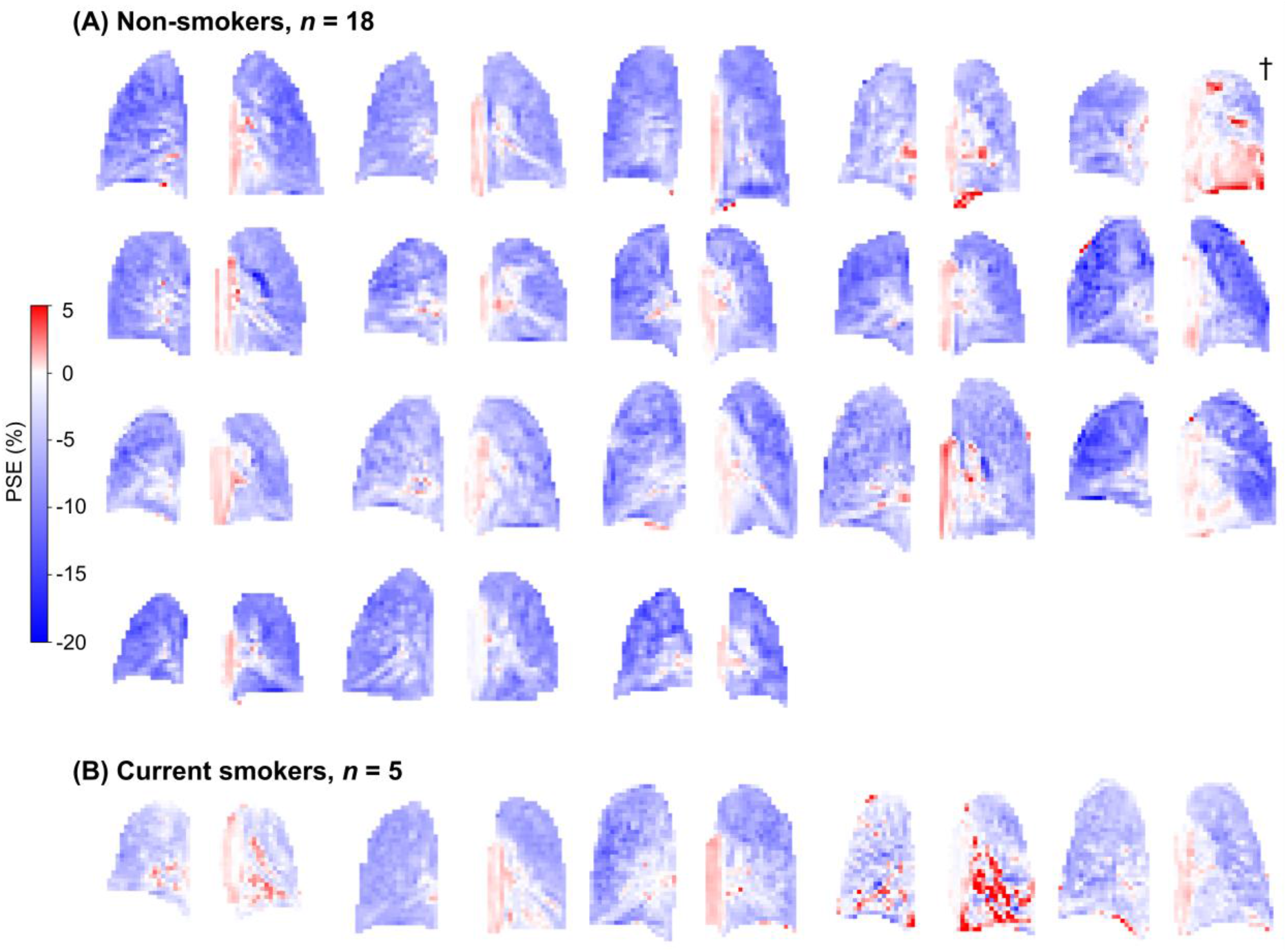
PSEICA maps of (A) non-smoker participants and (B) current smoker participants, all echo 1 data. The slice containing the aorta is shown for each participant; cardiac tissue is present in the lower left lung of the non-smoker labelled †. Minor variations between PSE_ICA_ maps arose due to differences in blood vessel and cardiac tissue content of the slice. An asymmetric color bar range of (-20, 5) is used to show the negative enhancement of the lung tissue and the positive enhancement of the heart and aorta (see also simulations in Supporting Information Figure S4).

Example ICA components extracted from each echo of a non-smoking participant are presented in Supporting Information Figures S6,7. For this participant, the Spearman correlation method identified the optimal OE ICA component in the run of ICA using 22 components for echo 1 and in the run of ICA using 23 components for echo 2. Figure 3 presents the mean echo 1 PSE time series within the lung mask for (A) PSE_MRI_ and (B) PSE_ICA_ of the same non-smoking participant; frequency spectra of the PSE time series are shown in Supporting Information Figure S8. The PSE_ICA_ time series exhibited well-defined cyclic enhancement compared to the PSE_MRI_ time series. The PSE_ICA_ frequency spectrum contained a sharp peak at the gas cycling frequency and minimal amplitudes at higher frequencies. Whereas the amplitude of the gas cycling frequency peak was lower in the PSE_MRI_ spectrum (spectra shown normalized by the maximum amplitude), and the PSE_MRI_ spectrum contained substantial frequency amplitudes within the cardiac and respiratory ranges.

**Figure 3:**
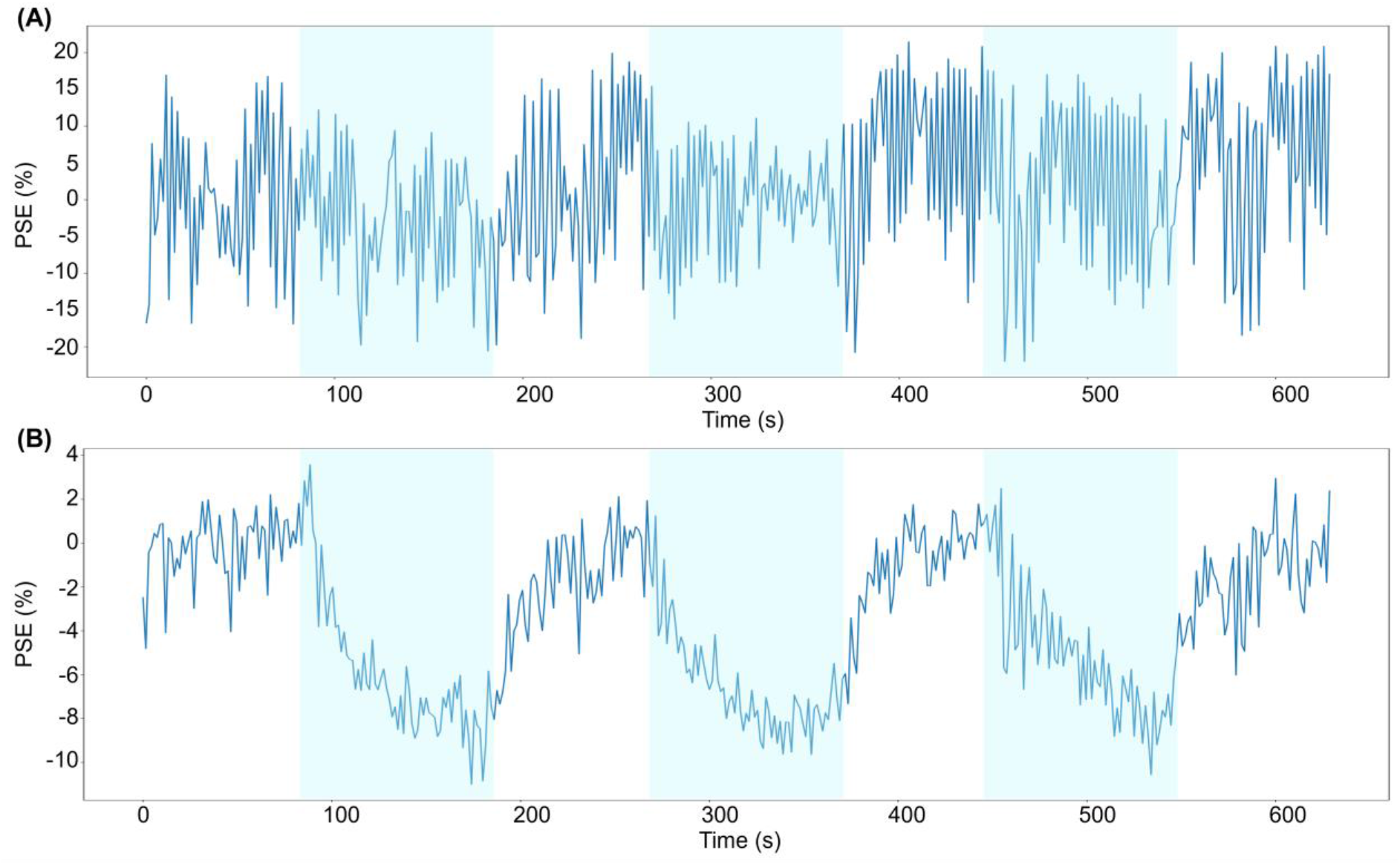
Time series of the mean PSE within the lung mask of (A) PSE_MRI_ data and (B) PSE_ICA_ data, both echo 1. Shaded blue periods indicate pure oxygen inhalation.

The signal enhancement of the lung tissue occurred with a negative PSE whereas the heart and aorta had a positive PSE, as demonstrated by the PSE_ICA_ maps in Figure 4(C). Regions of strong positive signal enhancement within the lung were observed in the PSE_MRI_ maps (Figure 4(B)) which were not seen in the PSE_ICA_ maps. The low amplitude opposite enhancements of the lung tissue from oxygenated blood contained within the heart and aorta were predicted by sequence-specific signal simulations^14, 25^ (Supporting Information Figure S4). Additionally, the simulations predicted a negative lung PSE at echo times greater than 0.23 ms due to the dominance of ΔT * effects – a negative lung PSE was observed by our experiment using echo times of 0.71 ms and 1.2 ms. For echo times shorter than 0.23 ms, the simulations predicted a small positive lung PSE due to the dominance of ΔT_1_ effects.

**Figure 4:**
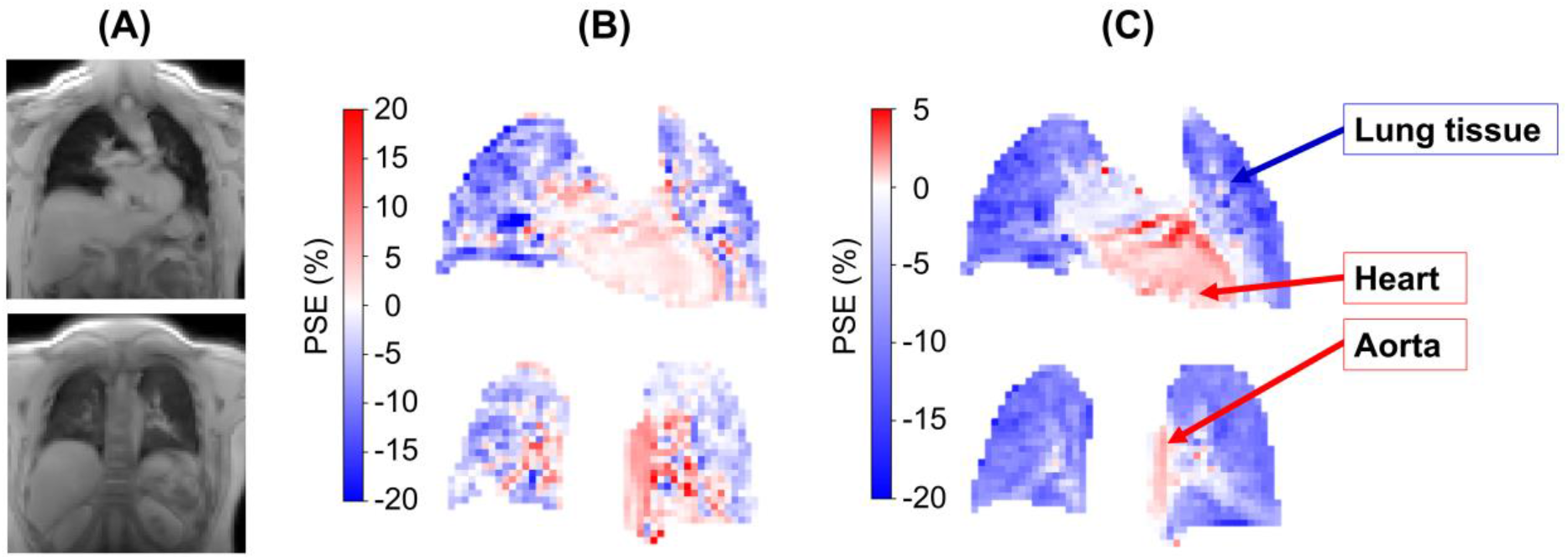
PSE maps of the non-smoking participant in Figure 3, all echo 1 data within the thoracic mask. (A) MRI images for anatomical reference; (B) PSE_MRI_ maps; (C) PSE_ICA_ maps. The signal enhancement of the lung tissue occurred with a negative PSE whereas the heart and aorta had a positive PSE. The PSE maps are presented using different color bar ranges to illustrate the different magnitudes of positive PSE. An asymmetric color bar range of (-20, 5) is used for the PSE_ICA_ maps (C) to show the negative enhancement of the lung tissue and the small positive enhancement of the heart and aorta (see also simulations in Supporting Information Figure S4). Whereas a symmetric color bar range of (-20, 20) is used for the PSE_MRI_ maps to display the regions of strong positive signal enhancement within the lung that were not observed in the PSE_ICA_ maps.

### 4.2. Smoking status PSE comparison

The PSE_ICA_ maps of all current smoker participants are displayed alongside those of non-smoker participants in Figure 2, with corresponding lung PSE_ICA_ time series shown in Supporting Information Figure S5. Figure 5 presents a comparison between the median lung PSE of current smoker and non-smoker participants. The median lung PSE_ICA_ was significantly smaller in current smokers than non-smokers: *p* = 0.002 for echo 1; *p* < 0.001 for echo 2. Whereas no significant difference was observed for the median lung PSE_MRI_: *p* = 0.154 for echo 1; *p* = 0.091 for echo 2.

**Figure 5:**
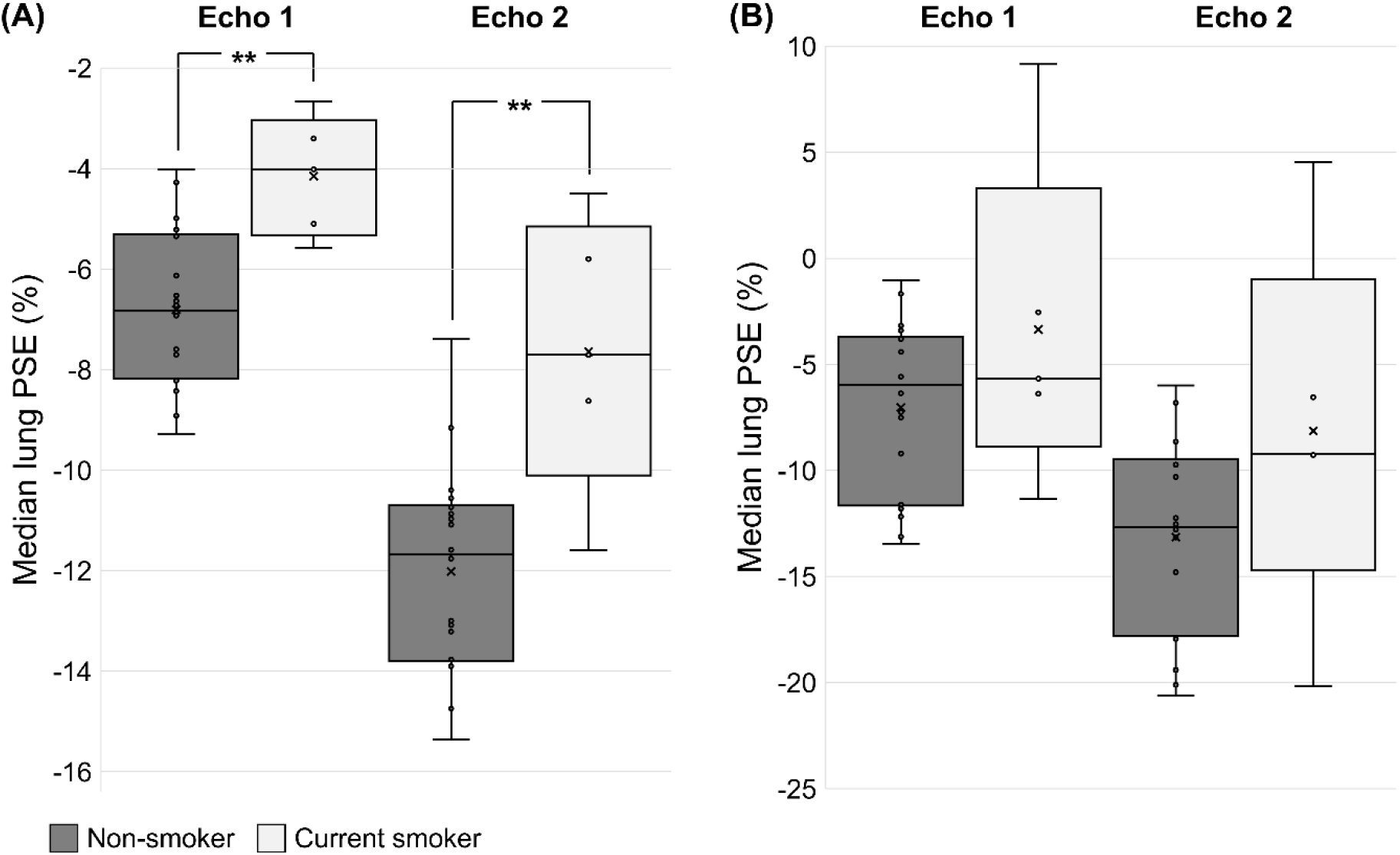
Comparison between the median lung PSE map value of non-smoker (gray) and current smoker (white) participants for each echo of (A) the PSE_ICA_ data and (B) the PSE_MRI_ data, without adjustment for the confounds of age and gender. The PSE_ICA_ of current smokers was significantly smaller than non-smokers for both echoes: *p* = 0.002 for echo 1; *p* < 0.001 for echo 2. No significant difference was present in the PSE_MRI_ data: *p* = 0.154 for echo 1; *p* = 0.091 for echo 2.

Supporting Information Table S4 contains a summary of the multiple regression results adjusted for the effects of age and gender. After adjustment, current smoking status remained significant for both echoes of the PSE_ICA_ data (*p* < 0.050 for echo 1; *p* = 0.033 for echo 2). In addition, gender was significant for PSE_ICA_ echo 2 (*p* = 0.043). For the PSE_MRI_ data, current smoking status remained non-significant after adjustment. Age was significant for both echoes of the PSE_MRI_ data (*p* = 0.034 for echo 1; *p* = 0.031 for echo 2).

### 4.3. Scan-rescan repeatability, ICA pipeline repeatability, and multi-site reproducibility

The scan-rescan repeatability, ICA pipeline repeatability, and multi-site reproducibility results are presented in Figure 6 (Bland-Altman plots) and Supporting Information Table S5 (statistical analysis). The repeatability analysis was presented in part at the Joint Annual Meeting ISMRM-ISMRT, London, 2023.^26^ Example scan-rescan PSE_ICA_ maps are displayed in Figure 7 for a non-smoker participant, in which similar spatial distributions of PSE were observed between scans. No significant biases were present in the median lung PSE_ICA_ value between scans: -0.085% for echo 1 and -0.023% for echo 2. The ICC values indicated good repeatability for both echoes: 0.807 for echo 1 and 0.907 for echo 2.

**Figure 6:**
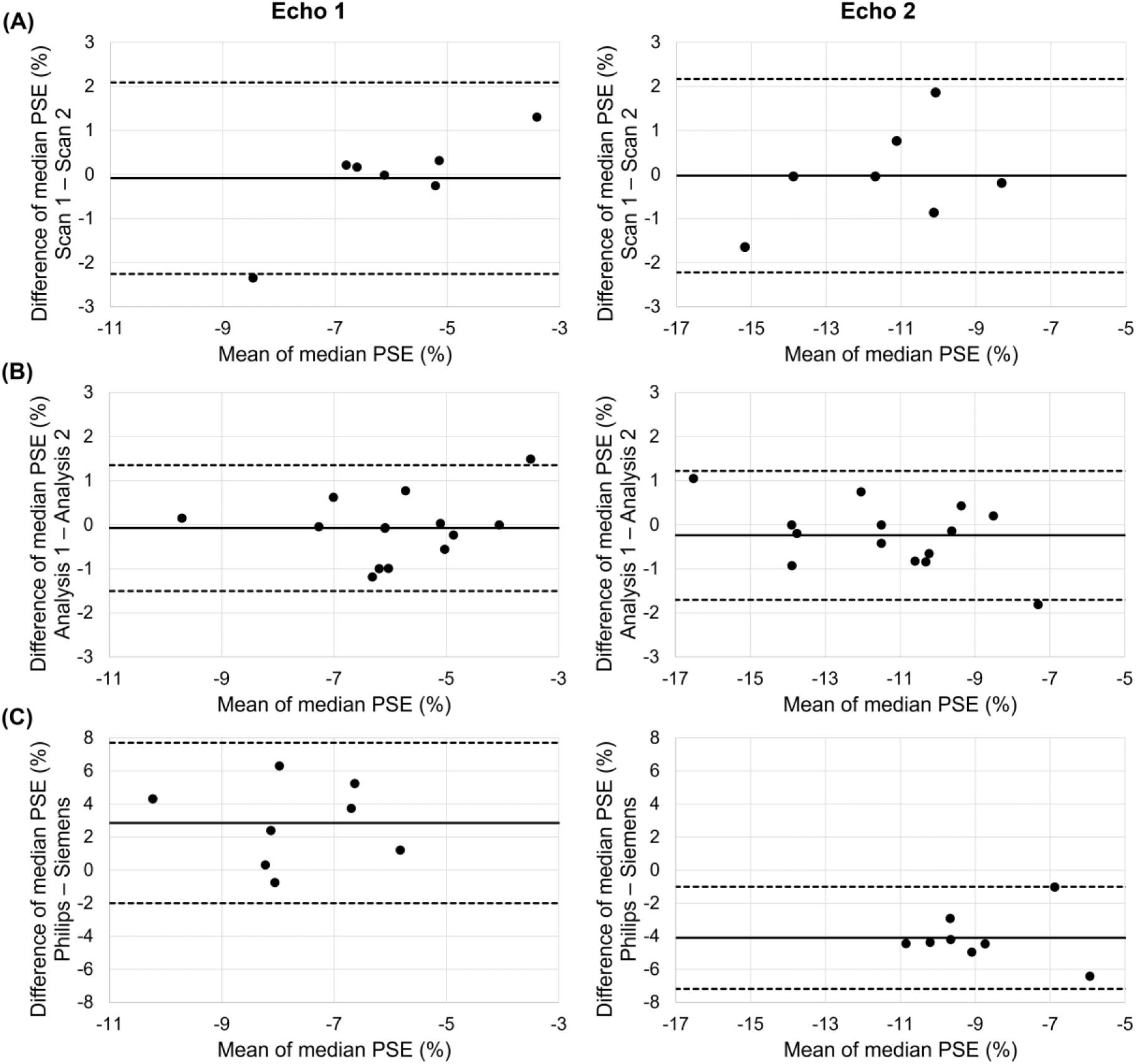
Bland-Altman plots of the median lung PSE_ICA_ values from the analysis of: (A) the scan-rescan repeatability; (B) the repeatability of ICA within the pipeline; and (C) the multi-site reproducibility. The solid black line indicates the bias and the dashed black lines indicate the limits of agreement. Bland-Altman analysis results are presented in Supporting Information Table S5.

**Figure 7:**
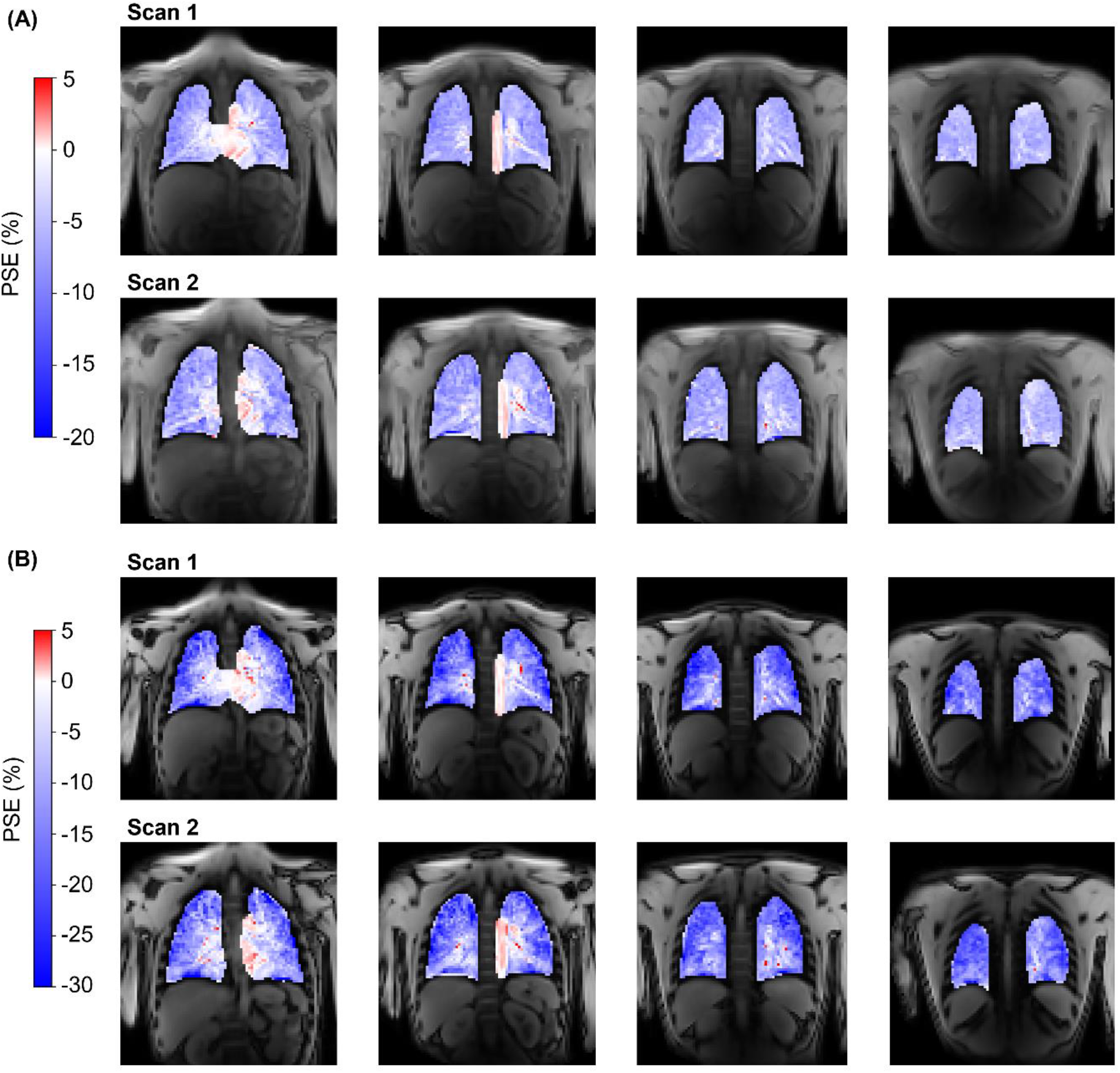
Example scan-rescan PSE_ICA_ maps for a non-smoker repeatability study participant. (A) echo 1 data from both scans; (B) echo 2 data from both scans. The lung PSE from echo 2 was more negative than echo 1 for both scans. Asymmetric color bar ranges of (-20, 5) for echo 1 and (-30, 5) for echo 2 are used to show the negative enhancement of the lung tissue and the positive enhancement of the heart and aorta (see also simulations in Supporting Information Figure S4).

Example PSE_ICA_ maps produced by the repeat application of the pipeline are displayed in Figure 8 for a non-smoker participant. Structures within the lung appeared consistent in the PSE_ICA_ maps from the repeated pipeline application to echo 1 data. Identical OE ICA components were extracted by the repeat pipeline application to echo 2 data. The repeat application of the ICA analysis pipeline displayed better repeatability than the scan-rescan analysis, demonstrated by the excellent ICC values of 0.926 for echo 1 and 0.958 for echo 2. No significant biases were observed: -0.075% for echo 1 and -0.240% for echo 2.

**Figure 8:**
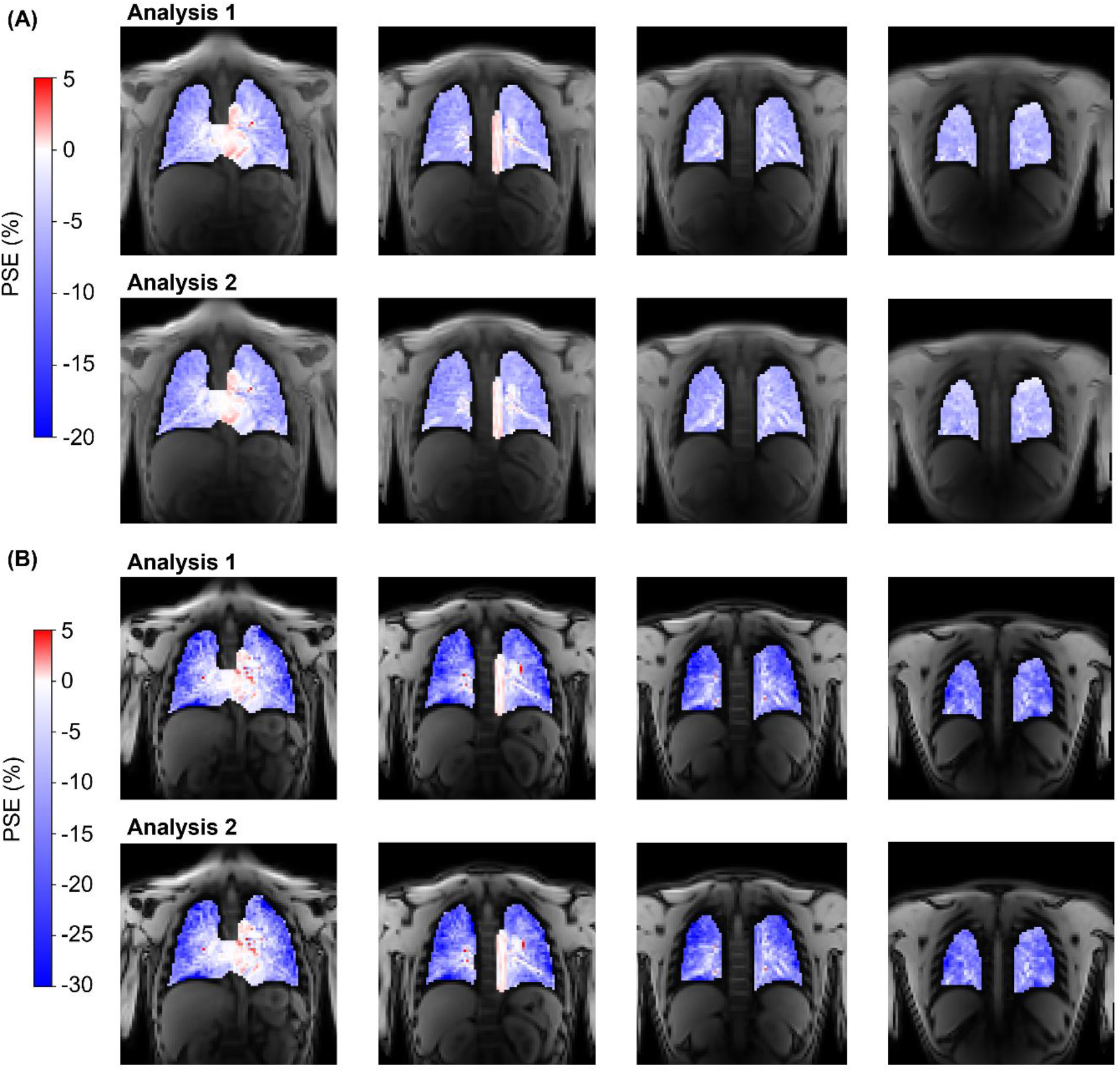
Example PSE_ICA_ maps produced by the repeat application of the pipeline for the non-smoker participant shown in Figure 7, for (A) echo 1 data and (B) echo 2 data. (Figure 7 presents the PSE_ICA_ scan-rescan maps; the current figure presents the PSE_ICA_ maps from the repeat pipeline application to data from scan 1.) The repeat pipeline application to echo 1 data produced similar PSE_ICA_ maps for which structures within the lung were consistent. Identical results were produced by the repeated pipeline application to echo 2 data. Asymmetric color bar ranges of (-20, 5) for echo 1 and (-30, 5) for echo 2 are used to show the negative enhancement of the lung tissue and the positive enhancement of the heart and aorta (see also simulations in Supporting Information Figure S4).

Figure 9 presents the PSE_ICA_ maps from both scans of a non-smoker reproducibility study participant. Similar spatial patterns of PSE were seen between the Siemens and Philips scans, however the PSE_ICA_ maps from the Siemens echo 2 data generally appeared to contain substantial noise-like patterns of positive PSE within the lung. Minor biases of 2.853% for echo 1 and -4.095% for echo 2, were observed. The signal simulation predicted a more negative PSE at each Siemens echo time than the corresponding Philips echo time, which would give rise to a positive bias between the two systems (Supporting Information Figure S4(B)). The predicted positive bias was observed for echo 1. However, contrary to the signal simulations, a negative bias was observed for echo 2 as the Siemens PSE_ICA_ was less negative than the Philips PSE_ICA_.

**Figure 9:**
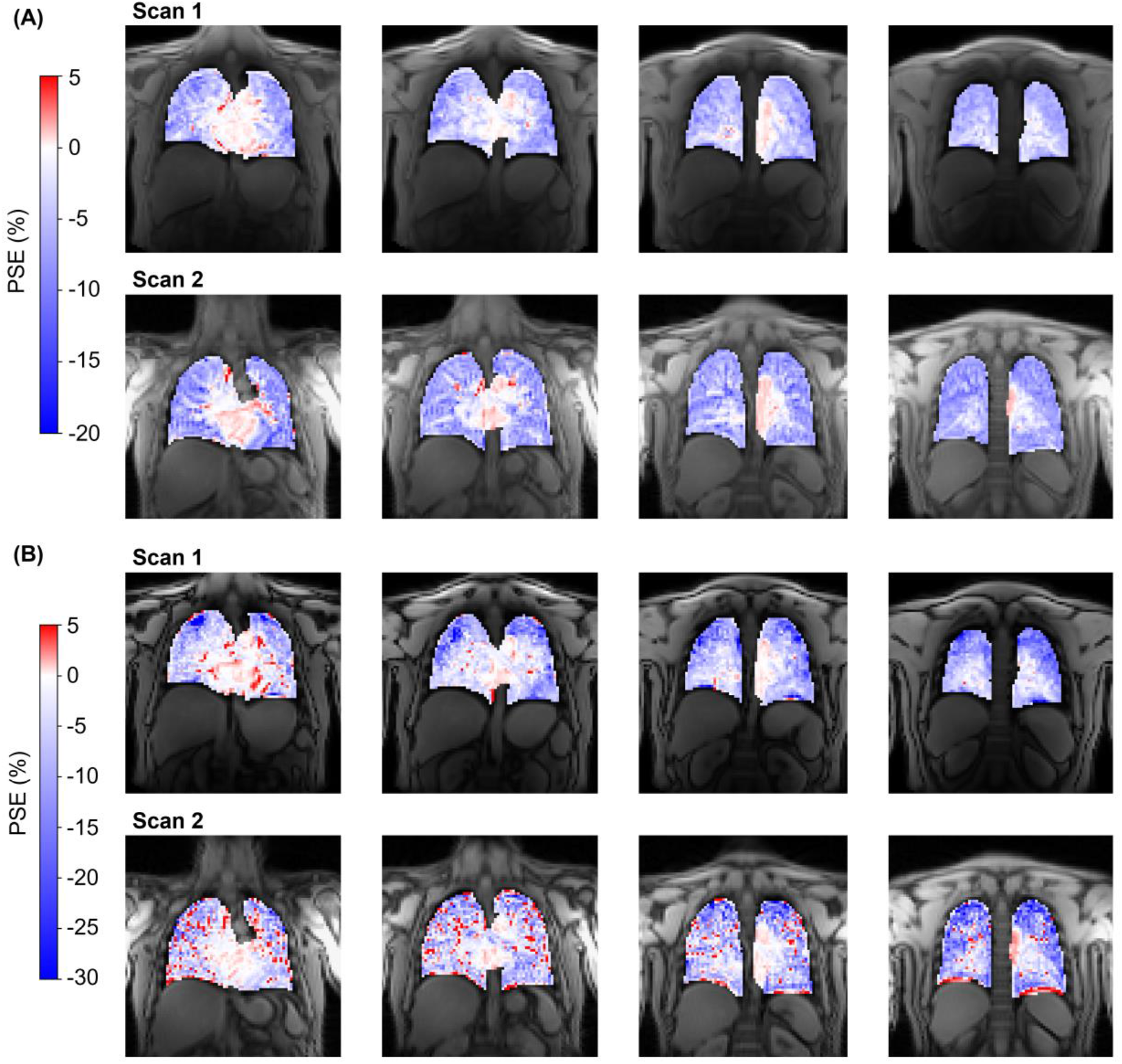
Example PSE_ICA_ maps for a non-smoker reproducibility study participant. (A) echo 1 data from both scans; (B) echo 2 data from both scans. Scan 1 was performed on a Philips Ingenia (TE_1_/TE_2_ = 0.71/1.2 ms) and scan 2 was performed on a Siemens MAGNETOM Vida (TE_1_/TE_2_ = 0.81/1.51 ms). Asymmetric color bar ranges of (-20, 5) for echo 1 and (-30, 5) for echo 2 are used to show the negative enhancement of the lung tissue and the positive enhancement of the heart and aorta (see also simulations in Supporting Information Figure S4).

## 5. Discussion

A previous study reported the use of ICA to extract the oxygen-induced signal change in pre- clinical tumor models.^8^ However, there are no published studies demonstrating the use of ICA to extract the oxygen-induced signal change in the human lung. To the best of our knowledge, we present the first application of ICA to lung OE-MRI. We developed an automatic analysis pipeline, utilizing ICA and a cyclic gas challenge, to extract the oxygen-induced signal change from the confounding factors present in lung MRI. The developed pipeline orders the ICA components using cross-correlation to identify the “optimal” oxygen- enhancement ICA component for each dataset. The objective and automatic approach for identifying the oxygen-enhancement signal increases the robustness of the pipeline and overcomes the main ambiguities of ICA in the ordering of components and in the number of components to use. We also show that our approach can be applied at 3 T to exploit the largely T *-driven contrast observed at this field strength.

### 5.1. Sensitivity to smoking status

The significant difference between the median lung PSE_ICA_ of current smokers and non-smokers, which remained after adjustment for the effects of age and gender, suggests that our OE-MRI acquisition and analysis method is sensitive to smoking-related changes of the lung. In contrast, no significant difference was observed between the median lung PSE_MRI_ of current smokers and non-smokers, both with and without adjustment for age and gender.

The lack of sensitivity to smoking status by the raw motion-corrected MRI data may be due to the presence of noise and confounding factors in the lung MRI data, which were reduced by ICA in the OE ICA component.

Age was significant for both echoes of the PSE_MRI_ data, but not for the PSE_ICA_ data. Age-related anatomical or physiological changes may have altered the MR signal, the effects of which ICA was able to reduce in the OE ICA component. However, the presented smoking status comparison is limited by the small number of participants in the study and by the gender imbalance – all current smoker participants were male. A larger study with balanced groups may provide further insights into the effects of age and gender on the oxygen-induced signal response of the lung.

### 5.2. Scan-rescan repeatability, ICA pipeline repeatability, and multi-site reproducibility

No significant PSE_ICA_ biases were observed for the scan-rescan study or for the repeat application of the pipeline to the scan-rescan data. The limits of agreement from the repeat application of the pipeline were smaller than, but of the same magnitude as, the scan-rescan study. This suggests that the algorithmic uncertainty of ICA contributes considerably to the overall variance of the OE-MRI experiment. The remaining variance in the scan-rescan study may be due to physiological variation including respiratory motion and motion-induced density changes, patient positioning, patient motion, and noise. Overall, the scan-rescan ICC values indicate a good repeatability of the OE-MRI ICA approach and suggest the pipeline is suitable for cross-sectional and longitudinal assessment.

When compared with the scan-rescan repeatability study, the reproducibility study biases were greater in magnitude, but still less than 5% (Supporting Information Table S5). Greater biases were expected for the reproducibility study due to the different echo times achievable on the two MR systems used. A negative bias was observed for echo 2 data as the Siemens PSE_ICA_ was less negative than the Philips PSE_ICA_, contrary to the signal simulations. The PSE_ICA_ map from Siemens echo 2, shown in Figure 9(B) [Scan 2], contained pixels with close to zero, and positive, PSE in the lung tissue. It is possible that the signal from the second echo of the Siemens scan was noise limited, which resulted in the less negative lung PSE_ICA_. Overall, we have demonstrated feasibility of the OE-MRI acquisition approach and analysis pipeline for use in multi-center studies and on different MRI vendor scanners.

### 5.3. Oxygen-induced MR contrast mechanisms

The lung tissue exhibited an opposite oxygen-induced signal change relative to the small positive PSE seen in the heart and aorta, which contain oxygenated blood. The opposite signal responses arise due to the dominance of T * contrast in lung tissue and T_1_ contrast in oxygenated blood, as predicted by the signal simulations shown in Supporting Information Figure S4. The oxygen-induced T * shortening of the lung, giving rise to a negative signal change, occurs due to an increase in the gas-tissue susceptibility gradients resulting from the elevated concentration of gaseous oxygen in the lung (contrast mechanism B in Figure 1).^9^

The oxygen-induced T_1_ contrast mechanism is generated by an increased concentration of dissolved oxygen in lung tissue water and blood, which causes T_1_ shortening (contrast mechanism A in Figure 1). This T_1_ shortening, and resulting positive signal change, contributes to the measured lung PSE due to the inherent T_1_-weighting in the spoiled gradient echo acquisition. However, as predicted by the signal simulations, the lung tissue signal change is dominated by ΔT * effects, instead of ΔT_1_ effects, at the echo times in our experiment, which gave rise to the negative PSE we observed in lung tissue.

For oxygenated blood in which gas-tissue susceptibility gradients do not exist, the dissolved oxygen-induced T_1_ contrast is predicted to dominate the OE-MRI signal change. This T_1_ contrast mechanism shortens the local T_1_ to generate a small positive PSE, as was observed in the aorta. The T * of oxygenated blood can be altered by susceptibility distortions generated by paramagnetic deoxyhemoglobin (BOLD contrast, mechanism C in Figure 1). The inhalation of pure oxygen can reduce the concentration of deoxyhemoglobin, causing T * to increase and generating a positive signal change. However, as oxygenated blood is almost fully saturated in healthy participants during air-inhalation,^12^ the inhalation of pure oxygen results in a minimal change to the concentration of deoxyhemoglobin. Hence, it is anticipated that BOLD contributions to ΔT * and the PSE of oxygenated blood are minor.^9, 13^

As discussed above, the T *-sensitive OE-MRI method we have developed at 3 T was sensitive to, and able to resolve, the different oxygen-induced MR signal changes of lung tissue from oxygenated blood, which has not previously been observed using OE-MRI. The majority of lung OE-MRI studies have focused on the use of T1-weighted sequences at 1.5 T^27^ due to the shorter T * of the lung and the reduced T_1_ relaxivity of oxygen^5^ at higher field strengths. Lung OE-MRI studies reported at 3 T have also utilized T_1_-weighted sequences, and as a result, did not exhibit sensitivity to the opposite T * and T signal changes of the lung and oxygenated blood.^28–30^ Hence, the use of T *-sensitive sequences is a promising direction for the use of OE-MRI at 3 T for functional lung imaging.

The measured MR signal is influenced by the changes in lung gas volume, tissue geometry, and lung proton density, that occur during the respiratory cycle. Changes in lung gas volume and tissue geometry alter T_2*_: lung T_2*_ is shorter at end-inspiration which decreases the MR signal intensity at this respiratory state.^4, 10, 31^ The MR signal intensity is further decreased at end-inspiration by the reduced proton density of the lung. These respiratory motion-induced changes to T_2*_ and the proton density were likely to have contributed to the noise within the lung observed in the PSE_MRI_ time series (Figure 3), and to have given rise to the substantial respiratory frequency amplitudes in the PSE_MRI_ frequency spectrum (Supporting Information Figure S8). In contrast, the amplitudes of respiratory frequencies were minimal in the PSE_ICA_ spectrum. Therefore, it is likely that ICA effectively removed signals relating to respiratory motion-induced T_2*_ and proton density changes from the OE ICA component.

The inhalation of pure oxygen can induce physiological changes which may contribute to the observed MR signal, such as changes to the pulmonary blood volume or blood vessel diameter. Some studies report no effect of hyperoxia on pulmonary blood volume,^32, 33^ whereas a recent OE-MRI study at 0.55 T by Wieslander et al.^34^ concluded that the pulmonary blood volume was altered by hyperoxia-induced vasodilation and resulted in a small but quantifiable contribution to the T_1_-weighted OE-MRI signal. Hyperoxia-induced vasodilation could alter our OE-MRI signal measurement in two main ways. Firstly, an increase in the pulmonary blood volume would increase the proton density of the lung, consequently increasing the measured lung MR signal. The increased lung MR signal would reduce the amplitude of the negative lung PSE, which could incorrectly be attributed to a reduced oxygen-enhancement response. Alternatively, T * could be altered by vasodilation and geometrical changes of the lung structure. Models by Weisskoff et al.^35^ and Boxerman et al.^36^ for susceptibility-induced ΔR * by vasculature in the brain suggest vasodilation causes T * to shorten. Vasodilation-induced T * shortening would augment the susceptibility related T * shortening that already occurs due to the inhalation of pure oxygen. Greater T * shortening would produce a more negative PSE, reinforcing the oxygen-induced signal changes within the lung tissue. The magnitude of such T * shortening is, however, difficult to predict for the greater magnetic susceptibility gradients within the lung relative to those used in the cerebral hemodynamics models by Weisskoff and Boxerman.

## 6. Conclusion

The novel application of ICA to lung OE-MRI enabled the separation of the ICA component relating to the oxygen-induced signal change from confounding factors present in the lung, improving the accuracy of lung OE-MRI analysis. At 3 T, the OE-MRI analysis pipeline we have developed was sensitive to, and enabled the resolution of, lung tissue from oxygenated blood. Our results demonstrated good scan-rescan and ICA pipeline repeatability, indicating robustness of the developed pipeline. We showed that the analysis pipeline was sensitive to smoking status, suggesting a likely sensitivity to pathology to be explored in future clinical studies.

## Acknowledgements

This work is supported by the EPSRC-funded UCL Centre for Doctoral Training in Medical Imaging (EP/L016478/1), the Cancer Research UK National Cancer Imaging Translational Accelerator (NCITA) award C1519/A28682 (UCL) and C19221/A28683 (University of Manchester), and Innovate UK award 104629. JM acknowledges funding from CRUK via the Network Accelerator Award Grant (A21993) to the ART-NET consortium and the Wellcome/EPSRC Centre for Interventional and Surgical Sciences (WEISS) (203145/Z/16/Z). This study represents independent research supported by the Manchester NIHR Biomedical Research Centre and by the National Institute for Health Research (NIHR) Biomedical Research Centre at The Royal Marsden NHS Foundation Trust and the Institute of Cancer Research, London. Many thanks to Lucy Caselton and Sumandeep Kaur for their help in acquiring the MR scans, to Dave Higgins (Philips) for his advice in developing the MR protocol, and to Gareth Ambler for statistical advice.

## Notes

### Competing Interest Statement

The authors have declared no competing interest.

