## Supplementary data for "Independent component analysis (ICA) applied to dynamic oxygen-enhanced MRI (OE-MRI) for robust functional lung imaging at 3 T"

### Supporting Information

Supporting Information Table S1: Details of (A) the non-smoker and (B) the current smoker groups involved in the healthy participant study.

| | | **(A)** | **(B)** |
|---|---|---|---|
| Participants | Total | 18 | 5 |
| | Male | 8 | 5 |
| | Female | 10 | 0 |
| Age (years) | Mean | 33 | 39 |
| | Range | 22-54 | 27-55 |
| Smoking status | Currently smoke regularly | 0 | 5 |
| Pack years | Mean | 1.2 | 6.4 |
| | Range | 0-8 | 1-14 |

Supporting Information Table S2: Details of the free-breathing dynamic lung OE-MRI sequences implemented on (A) the Philips Ingenia scanner and (B) the Siemens MAGNETOM Vida scanner.

| | **(A)** | **(B)** |
|---|---|---|
| Manufacturer | Philips | Siemens |
| Model | Ingenia | MAGNETOM Vida |
| Location | London | Manchester |
| Field strength (T) | 3 | 2.9 |
| RF coil used | 32-channel torso coil in combination with the posterior coil | 18 channel body coil in combination with the 32-channel spine coil |
| Max. gradient strength (mT/m) | 45 | 45 |
| Max. slew rate (T/m/s) | 200 | 200 |
| TR (ms) | 16 | 16 |
| Echoes | full | half |
| Minimum achievable $TE_1$ (ms) | 0.71 | 0.81 |
| Minimum achievable $TE_2$ (ms) | 1.2 | 1.51 |
| FOV (mm x mm) | 450 x 450 | 450 x 450 |
| No of slices | 4 | 4 |
| Slice thickness (mm) | 10 | 10 |
| Gap (mm) | 4 | 4 |
| Acquired matrix | 96 x 96 | 96 x 96 |
| Orientation | Coronal | Coronal |
| Pixel size (mm x mm) | 4.7 x 4.7 | 4.7 x 4.7 |
| Flip Angle (°) | 5 | 5 |
| Bandwidth (Hz/Px) | 4488 | 2000 |
| Parallel Imaging | N | N |
| NSA | 1 | 1 |
| Time resolution (s) | 1.54 | 1.54 |
| Number of dynamics | 420 | 420 |

Supporting Information Table S3: Details of the NiftyReg[1] parameters used to motion correct the dynamic images.

| | |
|---|---|
| Multi-resolution levels | 3 |
| Control point grid spacing | 3 voxels, 14.1 mm final spacing |
| Similarity metric | Locally normalized cross-correlation |
| Penalty terms and weights | Bending energy, 0.005; linear elastic energy, 0.01 |
| Transformation parameterization | Stationary velocity field |
| Maximum number of iterations per resolution level | 600, 300, 150 |

Supporting Information Table S4: Variable coefficients and their significance in the multivariable models generated to adjust for the confounds of age and gender on the comparison between the median lung PSE of non-smoker and current smoker participants. The median lung PSE of participants are plotted against age and gender in Supporting Information Figures S9 and S10, respectively.

Separate multiple regression models were created for: (A) echo 1 $PSE_{ICA}$ ($R^2$ = 0.474); (B) echo 2 $PSE_{ICA}$ ($R^2$ = 0.579); (C) echo 1 $PSE_{MRI}$ ($R^2$ = 0.301); (D) echo 2 $PSE_{MRI}$ ($R^2$ = 0.346). Current smoking status remained significant in the $PSE_{ICA}$ data for both echoes when adjusted for age and gender. Current smoking status was not significant for the $PSE_{MRI}$ data, both with and without adjustment for age and gender. Age was significant in the adjusted model for both echoes of the $PSE_{MRI}$ data.

| | **Variable** | **Standardized coefficient [95% confidence interval]** | | **Unstandardized coefficient [95% confidence interval]** | | **Significance** |
|---|---|---|---|---|---|---|
| **(A)** | Current smoking status | 0.409 | [0.000, 0.817] | 1.791 | [0.000, 3.582] | **0.050** |
| | Age | 0.205 | [-0.157, 0.568] | 0.040 | [-0.030, 0.109] | 0.250 |
| | Gender | 0.326 | [-0.072, 0.723] | 1.188 | [-0.261, 2.637] | 0.102 |
| **(B)** | Current smoking status | 0.404 | [0.037, 0.772] | 2.745 | [0.254, 5.236] | **0.033** |
| | Age | 0.294 | [-0.032, 0.620] | 0.088 | [-0.009, 0.185] | 0.074 |
| | Gender | 0.370 | [0.013, 0.727] | 2.090 | [0.074, 4.106] | **0.043** |
| **(C)** | Current smoking status | 0.100 | [-0.371, 0.571] | 1.199 | [-4.433, 6.830] | 0.661 |
| | Age | 0.455 | [0.037, 0.873] | 0.239 | [0.020, 0.458] | **0.034** |
| | Gender | 0.218 | [-0.240, 0.676] | 2.168 | [-2.390, 6.725] | 0.332 |
| **(D)** | Current smoking status | 0.133 | [-0.323, 0.589] | 1.848 | [-4.479, 8.174] | 0.548 |
| | Age | 0.451 | [0.046, 0.855] | 0.275 | [0.028, 0.521] | **0.031** |
| | Gender | 0.265 | [-0.178, 0.709] | 3.064 | [-2.056, 8.184] | 0.226 |

Supporting Information Table S5: Summary statistics from the analysis of: (A) the scan-rescan repeatability; (B) the ICA pipeline repeatability; and (C) the multi-site reproducibility. The bias, limits of agreement (LoA), repeatability coefficient (RC), and intra-class correlation coefficient (ICC), were calculated for the scan-rescan repeatability and ICA repeatability (A and B, respectively). Only the bias and LoA were calculated for the reproducibility study (C) as the measurement conditions were not identical for the two scans - longer echo times were implemented on the Siemens MAGNETOM Vida than the Philips Ingenia.

| | | **Bias (PSE %)** | **Limits of agreement (PSE %)** | **Repeatability coefficient (%)** | **ICC** |
|---|---|---|---|---|---|
| **(A)** | Echo 1 | -0.085 | [-2.257, 2.088] | 1.291 | 0.807 |
| | Echo 2 | -0.023 | [-2.218, 2.172] | 1.512 | 0.907 |
| **(B)** | Echo 1 | -0.075 | [-1.502, 1.352] | 1.008 | 0.926 |
| | Echo 2 | -0.240 | [-1.704, 1.223] | 1.152 | 0.958 |
| **(C)** | Echo 1 | 2.853 | [-1.992, 7.699] | | |
| | Echo 2 | -4.095 | [-7.180, -1.010] | | |

Supporting Information Figure S1: Diagram of the cyclic OE-MRI gas delivery scheme involving three periods of 100% $O_2$ inhalation. Gases were switched between medical air (21% $O_2$) and 100% $O_2$ every 1.5 minutes.

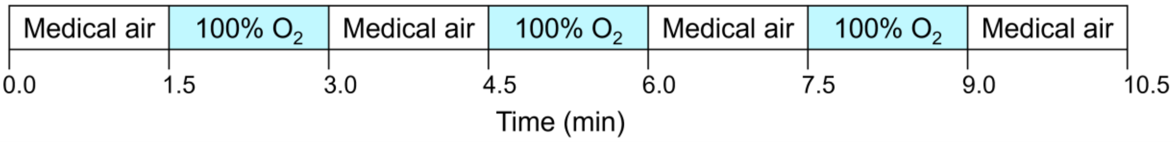


Supporting Information Figure S2: Example masks (light blue) overlaid on anatomical images. (A) lung mask: lung, excluding major vessels; (B) thoracic cavity mask: lung, heart, and major vessels.

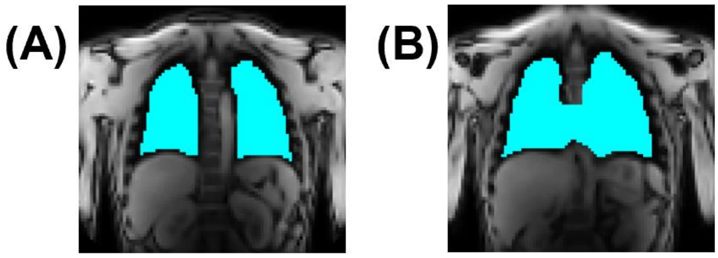

Supporting Information Figure S3: Diagram to illustrate the application of ICA to the dual-echo OE-MRI data and the approach devised to identify the optimal oxygen-enhancement ICA component.

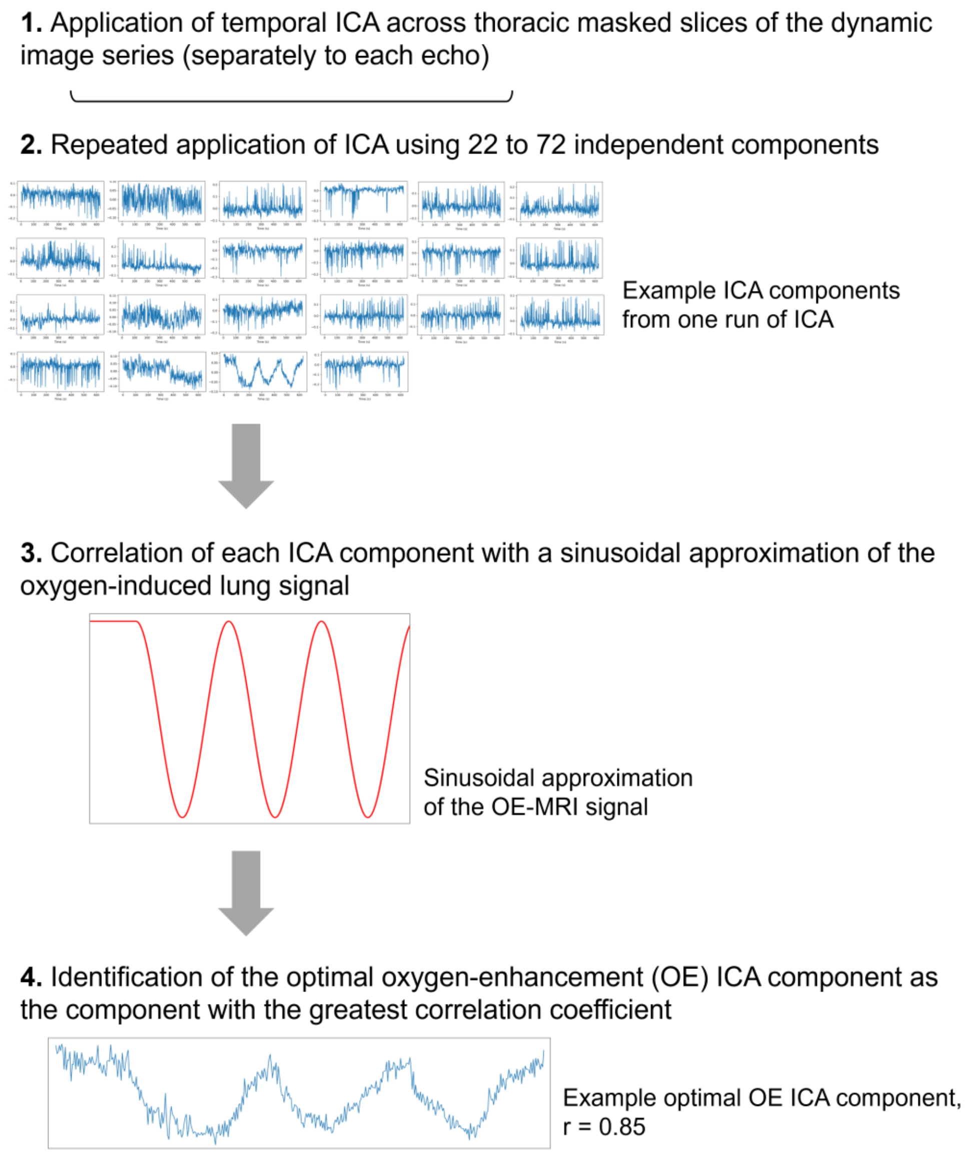

Supporting Information Figure S4: Sequence-specific MR simulations to predict the oxygen-enhancements of lung tissue and oxygenated blood for (A) the change in signal (ΔS) and (B) the percentage signal enhancement (PSE)[2,3]. Solid vertical lines indicate the echo times used for the Philips Ingenia ($TE_{1,P}$ and $TE_{2,P}$) and Siemens MAGNETOM Vida ($TE_{1,S}$ and $TE_{2,S}$) scans.

Lung relaxation times used: $T_{1,air}$ = 1281 ms and $T_{1,oxy}$ = 1102 ms[4]; $T_{2,air}^*$ = 0.68 ms and $T_{2,oxy}^*$ = 0.62 ms[2]. Blood (oxygenated) relaxation times used: $T_{1,air}$ = 1649 ms[5] and $T_{1,oxy}$ = 1354 ms[6]; $T_{2,air}^*$ = 59.4 ms and $T_{2,oxy}^*$ = 72.5 ms[7].

For TE < 0.23 ms (TE = 0.23 ms indicated by a dotted vertical line) the signal simulation predicts a positive lung PSE due to the dominance of $T_1$ effects, whereas for TE > 0.23 ms the simulation predicts a negative lung PSE due to the dominance of $T_2^*$ effects. Shown in (B), the PSE becomes more negative with increasing echo time.

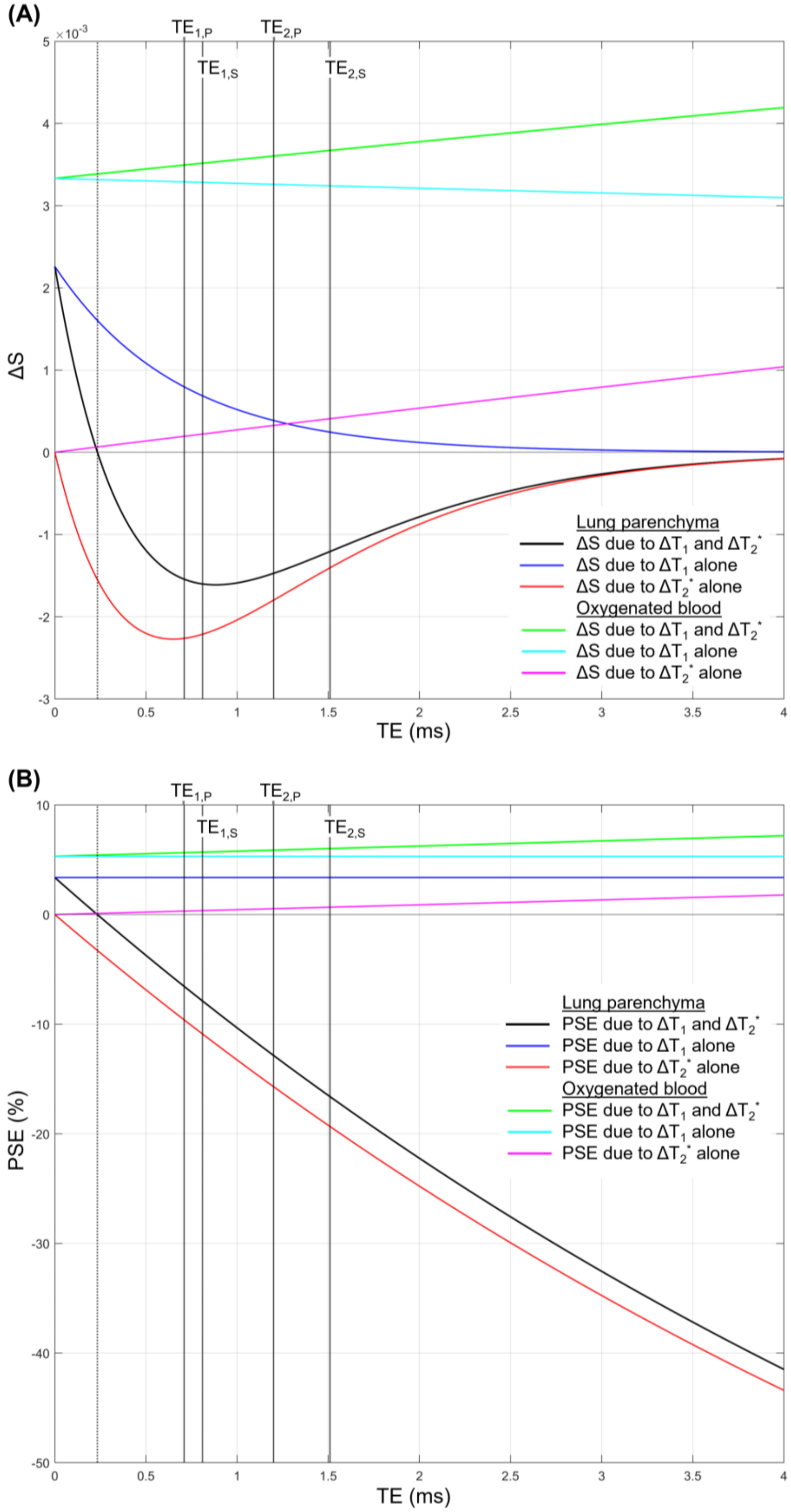
(A)
×10-3
TE1,P
TE2,P
TE1,S
TE2,S
ΔS
TE (ms)
Lung parenchyma
ΔS due to ΔT1 and ΔT2*
ΔS due to ΔT1 alone
ΔS due to ΔT2* alone
Oxygenated blood
ΔS due to ΔT1 and ΔT2*
ΔS due to ΔT1 alone
ΔS due to ΔT2* alone
(B)
PSE (%)
TE (ms)
Lung parenchyma
PSE due to ΔT1 and ΔT2*
PSE due to ΔT1 alone
PSE due to ΔT2* alone
Oxygenated blood
PSE due to ΔT1 and ΔT2*
PSE due to ΔT1 alone
PSE due to ΔT2* alone

Supporting Information Figure S5: The lung $PSE_{ICA}$ time series for the subjects presented in Figure 2. All echo 1 data for (A) non-smoker participants and (B) current smoker participants, shown with a y-axis range of -10% to 5% PSE.

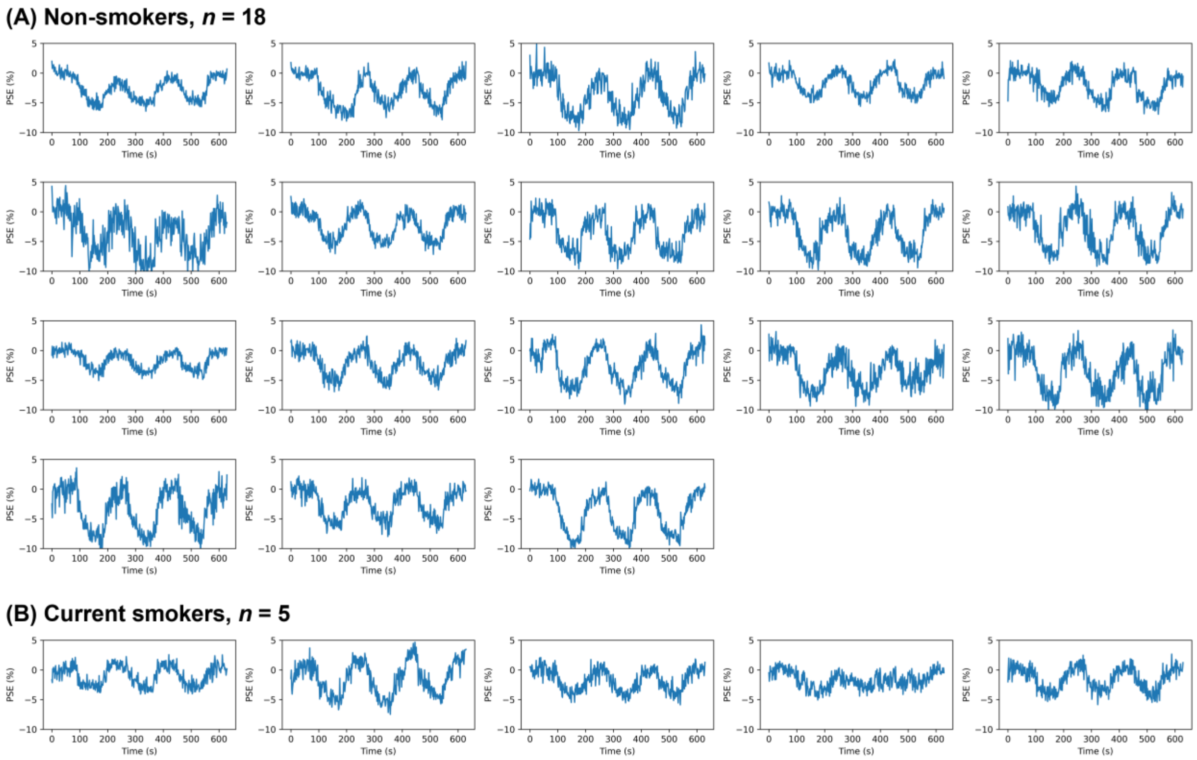

Supporting Information Figure S6: The ICA components extracted from echo 1 of a non-smoking participant, shown for the run of ICA in which the optimal OE ICA component was identified (22 components were used). The components are shown ordered by Spearman correlation value; the ordering metric value of each component is provided. The ordering approach successfully identified the OE ICA component as component 1. The OE ICA component displayed clear cyclic oxygen-enhancement with signal changes occurring upon the switching of gases. The ICA components have an arbitrary scaling and undetermined sign.

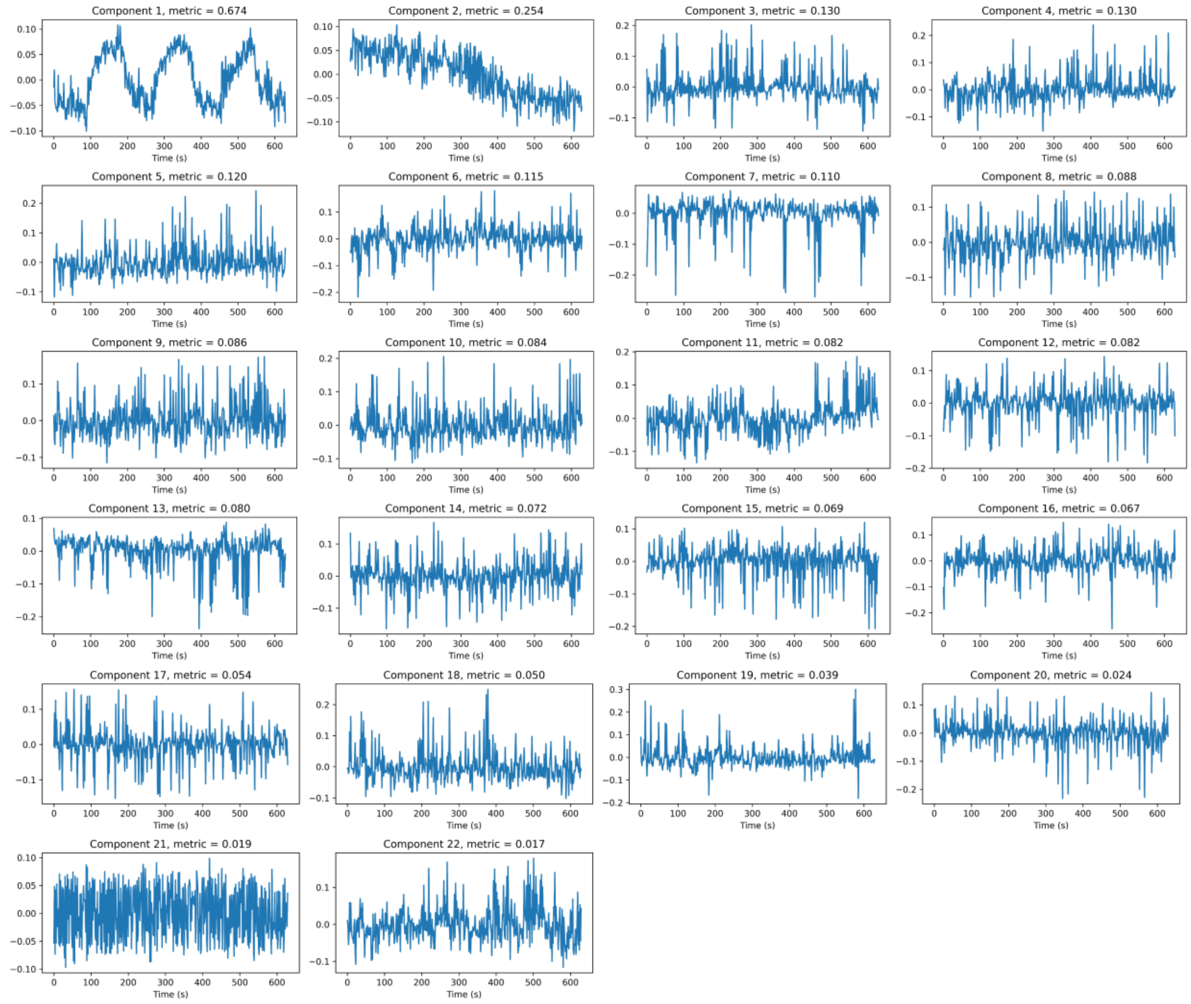

Supporting Information Figure S7: The ICA components extracted from echo 2 of the same non-smoking participant shown in Supporting Information Figure S6. The components presented are from the run of ICA in which the optimal OE ICA component was identified (23 components were used). The components are shown ordered by Spearman correlation value; the ordering metric value of each component is provided. The ordering approach successfully identified the OE ICA component as component 1. As for echo 1, the OE ICA component for echo 2 displayed clear cyclic oxygen-enhancement with signal changes occurring upon the switching of gases. The ICA components have an arbitrary scaling and undetermined sign.

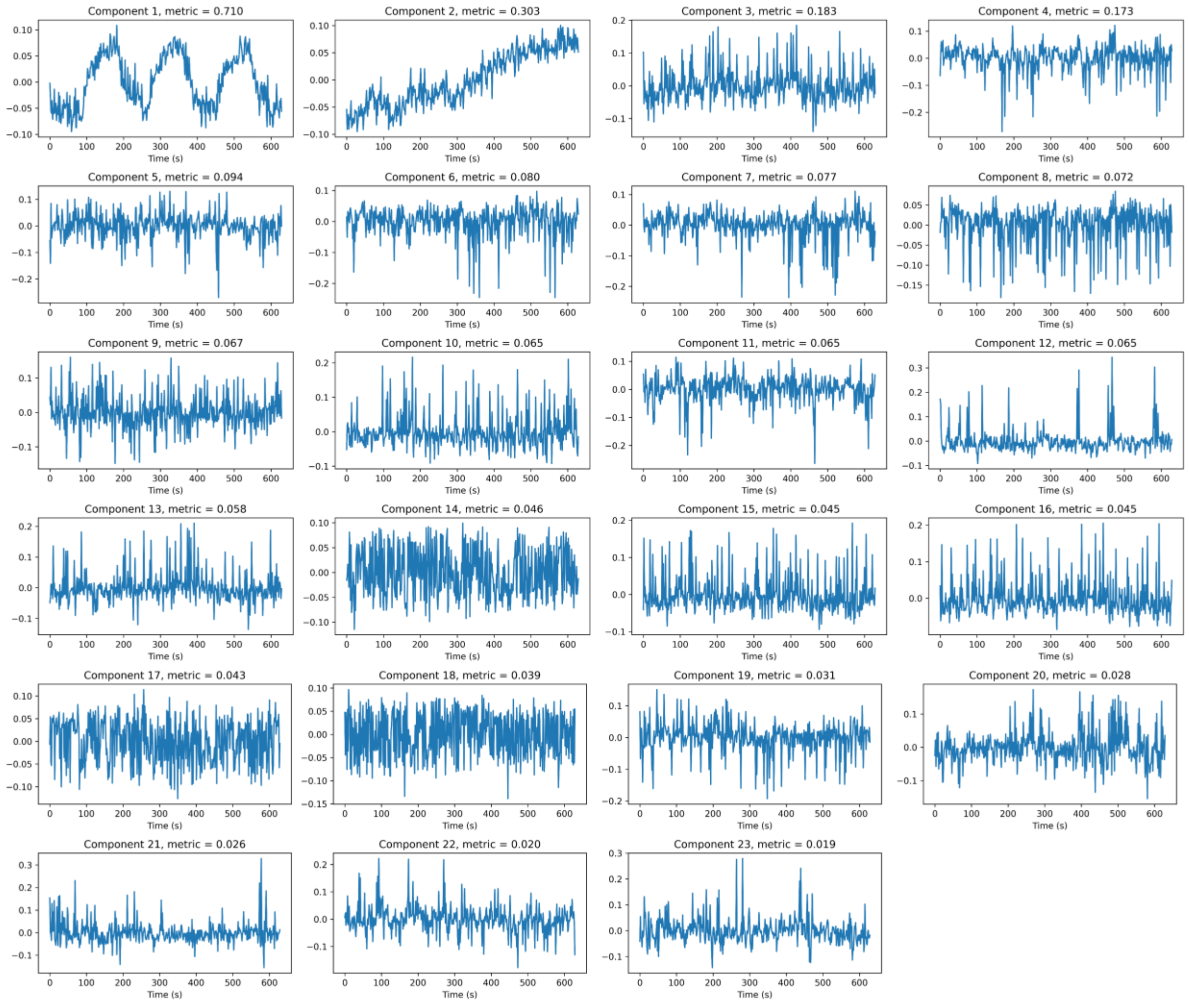

Supporting Information Figure S8: Frequency spectra of (A) the $PSE_{MRI}$ time series and (B) the $PSE_{ICA}$ time series shown in Figure 3. The frequency ranges associated with physiological motion and the OE-MRI gas cycling are indicated on the spectra: respiratory frequencies, $f_r$; aliased cardiac frequencies, $f_c$; and gas cycling frequency, $f_{OE}$ (also shaded in blue). The $PSE_{ICA}$ spectrum contained a peak at the $f_{OE}$ and minimal amplitudes at frequencies greater than $f_{OE}$. In contrast, the $PSE_{MRI}$ spectrum did not contain a sharp peak at $f_{OE}$ and displayed substantial frequency content above $f_{OE}$, particularly within $f_r$ and $f_c$.

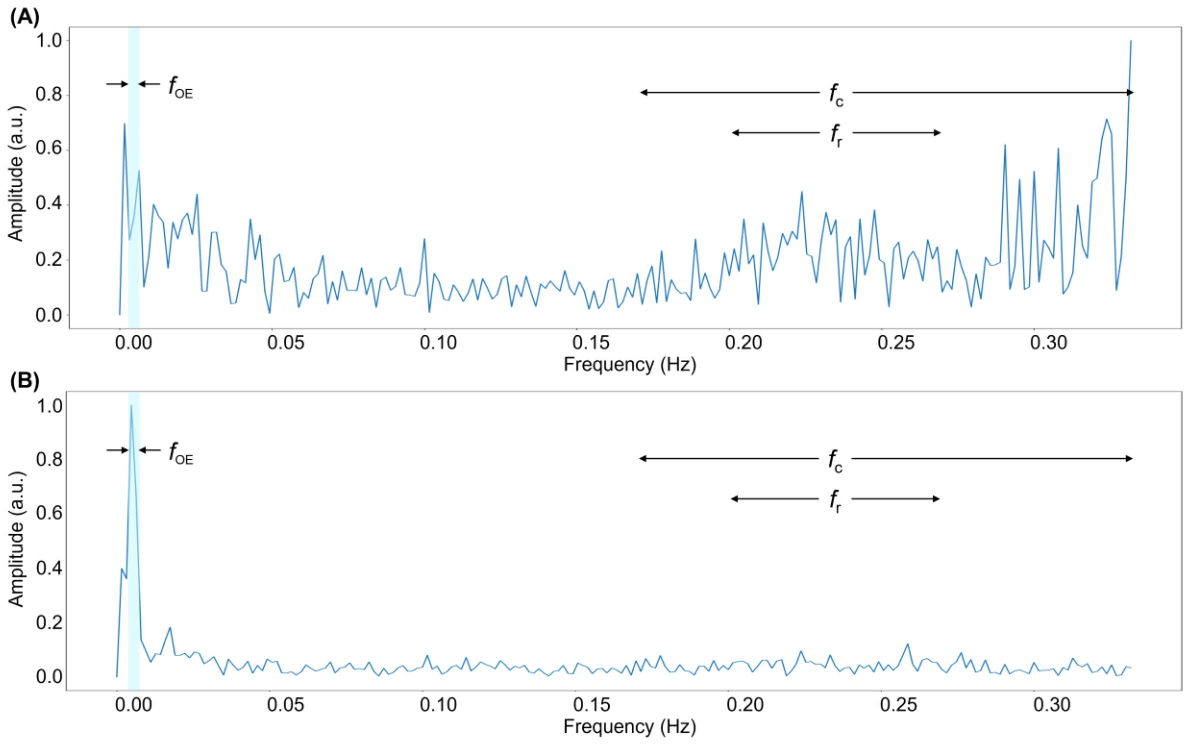

Supporting Information Figure S9: The median lung PSE map value of each participant plotted against age for each echo of (A) the $PSE_{ICA}$ data and (B) the $PSE_{MRI}$ data.

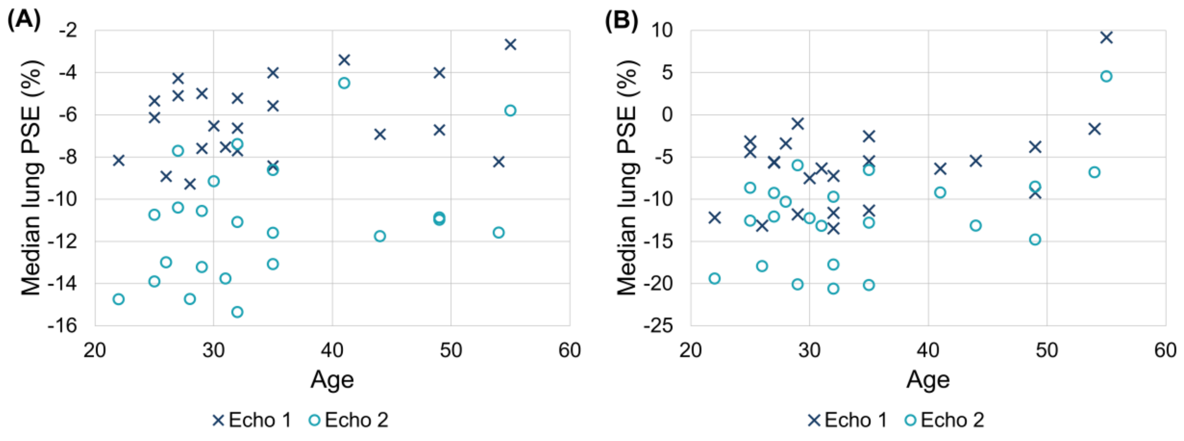


Supporting Information Figure S10: Comparison of the median lung PSE map value of male (gray) and female (white) participants for each echo of (A) the $PSE_{ICA}$ data and (B) the $PSE_{MRI}$ data.

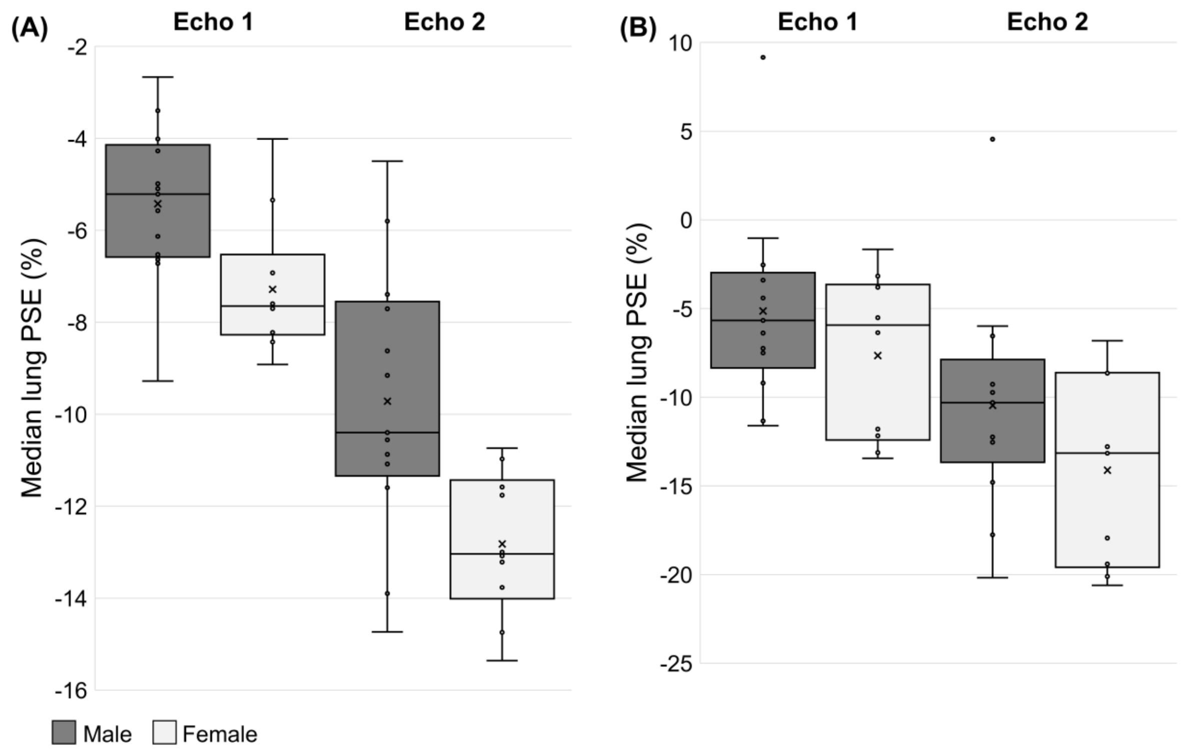
